# A novel role for the HLH protein Inhibitor of Differentiation 4 (ID4) in the DNA damage response in basal-like breast cancer

**DOI:** 10.1101/281196

**Authors:** Laura A. Baker, Christoph Krisp, Daniel Roden, Holly Holliday, Sunny Z. Wu, Simon Junankar, Aurelien A. Serandour, Hisham Mohammed, Radhika Nair, Chia-Ling Chan, Jessica Yang, Nicola Foreman, Breanna Fitzpatrick, Geetha Sankaranarayanan, Andrew M.K. Law, Chris Ormandy, Matthew J. Naylor, Andrea McFarland, Peter T. Simpson, Sunil Lakhani, Sandra O’Toole, Christina Selinger, Lyndal Anderson, Goli Samimi, Neville F. Hacker, Warren Kaplan, Jason S. Carroll, Mark Molloy, Alexander Swarbrick

**Affiliations:** The Kinghorn Cancer Center and Cancer Research Division, Garvan Institute of Medical Research, Darlinghurst, NSW 2010, Australia; St Vincent’s Clinical School, Faculty of Medicine, UNSW Sydney, NSW 2052, Australia; Australian Proteome Analysis Facility (APAF), Department of Chemistry and Biomolecular Sciences, Macquarie University, Sydney, Australia; Cancer Research UK, The University of Cambridge, Li Ka Shing Centre, Robinson Way, Cambridge CB2 0RE, UK; School of Medical Sciences and Bosch Institute, Sydney Medical School, The University of Sydney, New South Wales 2006, Australia; Centre for Clinical Research, Faculty of Medicine, The University of Queensland, Queensland, Australia; Pathology Queensland, The Royal Brisbane and Women’s Hospital, Herston, Brisbane, QLD, Australia; Department of Tissue Pathology and Diagnostic Oncology, Royal Prince Alfred Hospital, Camperdown, NSW 2050, Australia; Sydney Medical School, University of Sydney, NSW 2006, Australia; Rajiv Gandhi Centre for Biotechnology, Thycaud Post, Poojappura, Thiruvananthapuram-695014, Kerala, India; National Cancer Institute, National Institutes of Health, 9609 Medical Center Drive, Bethesda, MD 20892; School of Women’s and Children’s Health, University of New South Wales, and Gynaecological Cancer Centre, Royal Hospital for Women, Sydney, NSW, Australia

## Abstract

Basal-like breast cancer (BLBC) is a poorly characterised, heterogeneous disease. Patients are diagnosed with aggressive, high-grade tumours and often relapse with chemotherapy resistance. Detailed understanding of the molecular underpinnings of this disease is essential to the development of personalised therapeutic strategies. Inhibitor of Differentiation 4 (ID4) is a helix-loop-helix transcriptional regulator required for mammary gland development. ID4 is overexpressed in a subset of BLBC patients, associating with a stem-like poor prognosis phenotype, and is necessary for the growth of cell line models of BLBC, through unknown mechanisms. Here, we have defined a molecular mechanism of action for ID4 in BLBC and the related disease highgrade serous ovarian cancer (HGSOV), by combining RIME proteomic analysis and ChIP-Seq mapping of genomic binding sites. Remarkably, these studies have revealed novel interactions with DNA damage response proteins, in particular, mediator of DNA damage checkpoint protein 1 (MDC1). Through MDC1, ID4 interacts with other DNA repair proteins (γH2AX and BRCA1) at fragile chromatin sites. ID4 does not affect transcription at these sites, instead binding to chromatin following DNA damage and regulating DNA damage signalling. Clinical analysis demonstrates that ID4 is amplified and overexpressed at a higher frequency in *BRCA1*-mutant BLBC compared with sporadic BLBC, providing genetic evidence for an interaction between ID4 and DNA damage repair pathways. These data link the interactions of ID4 with MDC1 to DNA damage repair in the aetiology of BLBC and HGSOV.

## Introduction

Breast cancer is a heterogeneous disease. Gene expression signatures delineate five major subtypes with distinct pathologies, treatment strategies and clinical outcomes (Perou et al., 2000, Prat and Perou, 2011, Rouzier et al., 2005, Sørlie et al., 2001). The basal-like subtype (BLBC), accounting for ∼18% of diagnoses, is a subtype with particularly poor prognosis, largely due to the molecular and clinical heterogeneity of these tumours, and the corresponding lack of targeted therapeutics. The molecular drivers of BLBC are poorly understood, thus there are limited available targeted therapies.

A molecular driver of a subset of BLBCs is mutation in the breast and ovarian cancer susceptibility gene (*BRCA1*) (Miki et al., 1994, Turner and Reis-Filho, 2006, Turner et al., 2007). Mutations in *BRCA1* occur in approximately 0.25% of European women, predisposing them to breast and ovarian cancer, particularly to poor prognosis subtypes including BLBC and high-grade serous ovarian cancer (HGSOV) (Miki et al., 1994). These two subtypes of cancer are similar in terms of their gene expression profiles and genetic dependencies, and thus present similar sensitivity to therapeutic targeting (Network, 2012, Network, 2011, Marcotte et al., 2016). BRCA1 has many cellular functions including transcription and gene splicing, yet is best known for its role in mediating DNA damage repair (Mullan et al., 2006, Scully et al., 2004, Wu et al., 2010). BRCA1 coordinates efficient repair of double stranded DNA breaks through the Homologous Recombination (HR) pathway (Scully et al., 2004). In the absence of functional BRCA1, cells accumulate mutations, genomic instability and demonstrate an increased frequency of genomic rearrangements (Holstege et al., 2010, Vollebergh et al., 2010). BRCA1 mutations confer synthetic lethality to platinum based chemotherapies and PARP (poly ADP ribose polymerase) inhibitors, both of which target the DNA damage response (Audeh et al., 2010, Domagala et al., 2011, Tassone et al., 2009, Vollebergh et al., 2010).

To explore further molecular drivers of BLBC, we investigated the Helix-Loop-Helix (HLH) transcriptional regulator Inhibitor of differentiation 4 (ID4). We and others have previously shown ID4 to be a master regulator of mammary stem cell self-renewal in the normal mammary gland (Junankar et al., 2015, Best et al., 2014, Dong et al., 2011), as well as important for the aetiology of BLBC. ID4 is overexpressed in a subset of BLBC patients, marking patients with poor survival outcome, and is necessary for the growth of BLBC cell lines (Beger et al., 2001, Branham et al., 2016, Crippa et al., 2014, de Candia et al., 2006, Junankar et al., 2015, Rold á n et al., 2006, Shan et al., 2003, Thike et al., 2015, Wen et al., 2012). Precisely how ID4 mediates this function in BLBC is unclear.

ID proteins (ID1-4) lack a basic DNA binding domain, and thus, their classical mechanism of action is believed to entail dominant-negative regulation of canonical binding partners; basic HLH (bHLH) transcription factors. ID proteins dimerise with bHLH proteins and prevent them from interacting with DNA, affecting the transcription of lineage-specific genes (Benezra et al., 1990, Jen et al., 1992, Loveys et al., 1996, Langlands et al., 1997, Roberts et al., 2001). Yet this model of ID protein function is largely based on evidence from studies of ID1-3 in non-transformed fibroblasts, neural and embryonic tissue. ID proteins are tissue specific in their expression and function and hence this model may not apply to all four ID proteins across various tissues and in disease (Jen et al., 1996, Langlands et al., 1997, Melnikova et al., 1999, Norton, 2000, Chaudhary et al., 2001, Engel and Murre, 2001, Ruzinova and Benezra, 2003, Perk et al., 2005, Asirvatham et al., 2006). Indeed, contrary mechanisms have been described for ID2 in liver regeneration, with ID2 interacting with chromatin at the c-Myc promoter as part of a multi-protein complex to repress c-Myc gene expression (Ferrer-Vicens et al., 2014, Rodr í guez et al., 2006), and ID4 has been shown to bind to and suppress activity of the ERα promoter and regions upstream of the ERα and FOXA1 genes in mouse mammary epithelial cells (Best et al., 2014). These data suggest that despite lacking a DNA binding domain, ID proteins may interact with chromatin complexes under certain conditions. However, no studies have systematically mapped the protein or chromatin interactomes for any ID family member.

To this end, we applied chromatin immunoprecipitation-sequencing (ChIP-seq) to interrogate the ID4-chromatin binding sites, Rapid Immunoprecipitation and Mass spectrometry of Endogenous proteins (RIME) to determine the ID4 protein interactome. In addition, ID4 knockdown and RNA-sequencing analysis was used to determine transcriptional targets of ID4. Integrating these data, we have defined a novel mechanism of action for ID4 in regulating the DNA damage response.

## Results

### ID4 interacts with chromatin without regulating transcription

To investigate the molecular function of ID4, we first examined ID4 protein expression and cellular localisation across a panel of breast and ovarian cancer cell lines. Ovarian cancer cell lines were included due to the established molecular similarities between BLBC and HGSOV (Network, 2012, Network, 2011, Marcotte et al., 2016). Furthermore, evidence suggests ID4 may play an important role in the aetiology of both BLBC and HGSOV (Ren et al., 2012, Junankar et al., 2015). As described previously (Junankar et al., 2015), ID4 is predominantly expressed by cell lines of the BLBC subtype (Supplementary Figure 1a). Variable ID4 protein expression was observed across the ovarian cell lines, and unlike in BLBC ID4 did not associate specifically with HGSOV, the more aggressive ovarian cancer subtype. Four ID4 expressing models were selected for further analysis: MDA-MB-468 (BLBC), HCC70 (BLBC), HCC1954 (HER2-Enriched) and OVKATE (HGSOV), because these models represent different biological systems with similar levels of ID4 expression.

**Figure 1:**
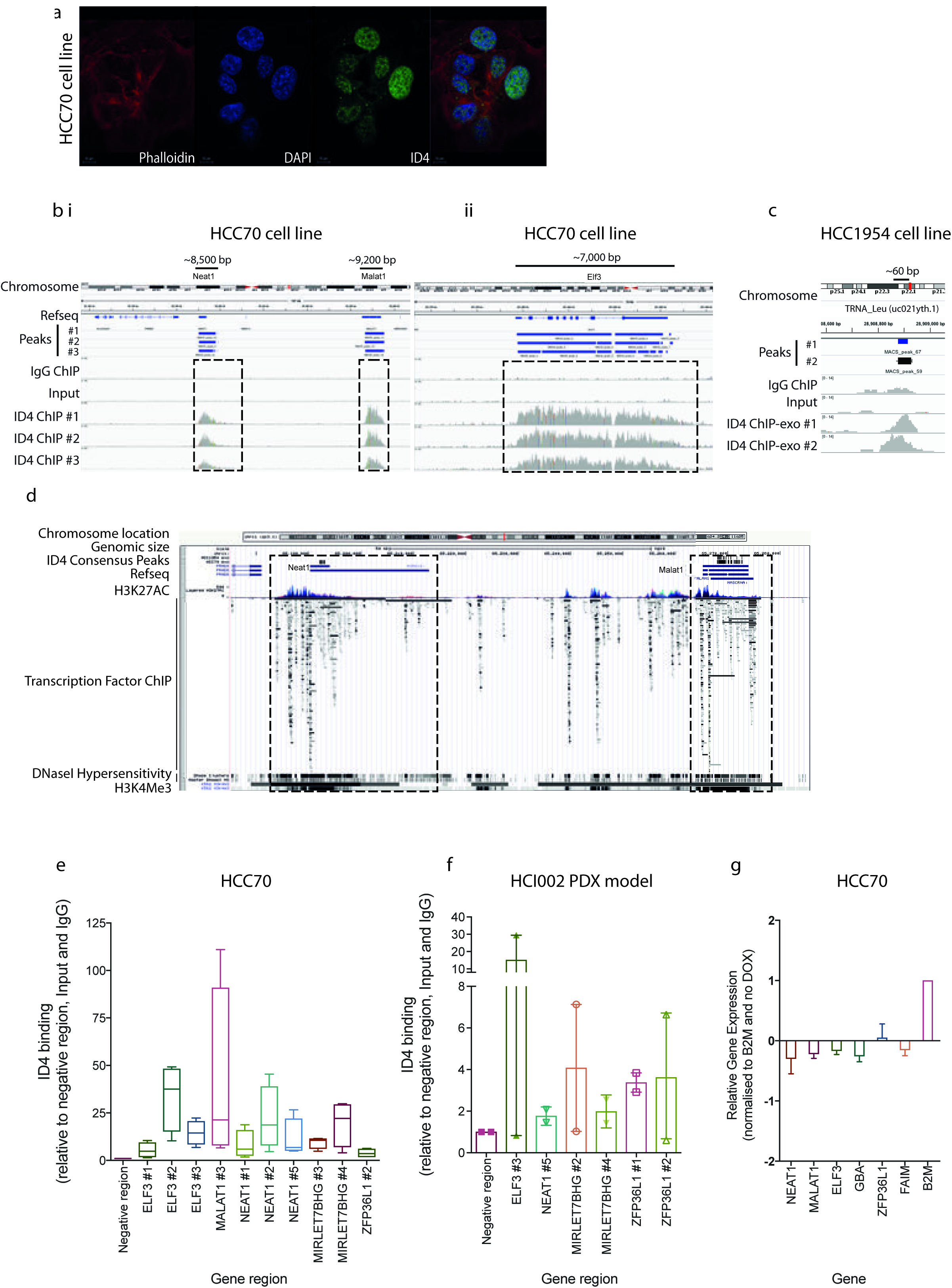
ID4 binds to chromatin in BLBC cell lines and PDX models and does not regulate transcription. A; Immunofluorescence analysis of ID4 protein expression in the HCC70 BLBC cell line, blue; nuclear marker DAPI, red; cytoskeletal marker phalloidin, green; ID4. Scale on images. B; Three technical replicates of ID4 ChIP-seq analysis in HCC70 cell line. IgG ChIP-seq and Input controls are shown for comparison. ID4 binding to i) the genomic region encoding the long non-coding RNAs NEAT1 and MALAT1 and ii) the protein coding gene ELF3. C; ID4 binding to tRNA_Leu (uc021yth.l) in HCC1954 cell line measured by ChIP-exo. Identification of ID4 binding enrichment identified by MACS peak-calling algorithm. Chromosome location, transcription start site (TSS) and Refseq information tracks displayed. Reads have been aligned to the human reference genome Hgl9 and peaks called using MACs peak calling algorithm (v2.0.9) (Serandour et al., 2013). Images contain ChIP-seq coverage data and the peaks called for each ID4 technical replicate and the consensus peaks called for all three ID4 ChIP-seq technical replicate for selected gene regions. ID4 binding is shown in comparison to IgG and Input data for the same region. Data visualised using IGV (Robinson et al., 2011, Thorvaldsdottir et al., 2013). Transcription Start Site (TSS) indicated with black arrow. D; Alignment of ID4 ChIP-exo peaks with DNAse hypersensitivity clusters, Transcription Factor ChIP and histone marks H3K27Ac and H3K4Me3 ChIP-seq at NEAT1 and MALAT1, UCSC Genome Browser. E; ID4 ChIP-qPCR analysis in HCC70 cells. Multiple primers were designed to tile across the large ID4 binding sites. ID4 binding normalised to input DNA and to a region not bound by ID4 (negative region) and represented as fold-change over IgG control. Ratio paired t-test, all p < 0.05. F; top: ID4 IHC in HCI002 triple-negative patient-derived xenograft model, bottom: ID4ChIP-qPCR analysis in HCI002 model. ID4 binding normalised as for E. G; qRT PCR analysis of mRNA transcript expression of ID4-bound genomic regions following depletion of ID4 in HCC70 cells using a lentiviral, doxycycline-inducible short hairpin RNA #2 (SMARTChoice). Data normalised to B2M housekeeping gene and to HCC70 cells treated with vehicle control. Un-paired t-test of control primer B2M with each primer set: NEAT1 p=0.017, MALAT1, ELF3, GBA and FAIM p<0.0001, ZFP36L1 p=0.008.

ID4 was shown to predominantly localise to chromatin, as evidenced by distinct punctate staining in the nucleus, with a proportion of ID4 staining overlapping with DAPI-low nuclear regions (Figure 1a), typical of uncondensed euchromatin. Due to this localisation, and previous association of ID4 with enhancer regions in normal mammary epithelial cells (Best et al., 2014), chromatin immunoprecipitation and sequencing (ChIP-seq) was used to interrogate the precise ID4-chromatin binding sites in BLBC. ID4 ChIP-seq was conducted on asynchronous HCC70 cells in three biological replicates and compared with IgG ChIP and an input control. Western blot analysis confirmed successful immunoprecipitation of ID4 (Supplementary Figure 1b). ChIP-seq data was integrated by removing signal identified in the input and IgG controls and examining the overlap between the three ID4 ChIP-seq replicates. This analysis revealed that ID4 associated with a seven reproducible sites across the genome (Supplementary Figure 2a and Supplementary Table 1). Some of these were focussed, small binding events typical of transcription factor binding; as examples, ID4 binding to the transcription start site of *FAIM* and *GBA* (140-230 bp) (Supplementary Figure 2c and d) was typical of transcription factor binding peaks. In other locations, ID4 binding was spread over very large regions of DNA, up to lOkb in length, which is reminiscent of some histone marks, and not transcription factor binding profiles. ID4 bound primarily to gene bodies of the protein-coding genes *ELF3 and ZFP36L1*, the long noncoding RNAs *NEAT1* and *MALAT1* and the microRNA host-gene *MIRLET7BHG.* The MACS peak calling algorithm identified multiple peaks per gene for these genes, due to wide spread binding across the region (*ELF3, NEAT1, MALAT1, MIRLET7BHG* and *ZFP36L1*) (Figure 1b and Supplementary Figure 2b-e). This binding is absent from intergenic regions (Figure 1b).

Due to the large expanses of ID4 binding, ChIP-exonuclease (ChIP-exo) was employed to provide higher resolution mapping of ID4 binding. ChIP followed by exonuclease digestion and DNA sequencing identifies precise transcription factor binding sites with high resolution than traditional ChIP-seq (Serandour et al., 2013). ID4 ChIP-exo was conducted in biological duplicate in HCC70 and HCC1954 breast cancer cell lines. Analysis of ChIP-exo data using the MACS peak calling algorithm identified significant enrichment of ID4 binding to the genomic regions encoding NEAT1, MALAT1 and GBA. Signal was observed at ELF3, MIRLET7BHG and ZFP36L1, however these regions were not identified through MACS analysis (Supplementary Figure 3). This method was unable to further resolve ID4-chromatin binding information and the data resembled the ChIP-seq data. This suggests that ID4 forms large contiguous and uninterrupted interactions with expanses of chromatin. Additional ID4 binding peaks were identified including small binding events (150-200 bp width) within the gene bodies of KDM4C (identified in HCC1954 cell line) and ERRFI1 (identified in HCC1954 and HCC70 cell lines), which was not identified in ChIP-seq on HCC70 cell line. Additionally, MACS analysis identified ID4-bound peaks in several transfer RNAs (tRNA) in the HCC1954 genome (Figure 1c). Signal was observed at tRNAs in HCC70 cells, however, these regions were not identified through MACS analysis.

ChIP-qPCR was used to validate ID4 binding in HCC70, HCC1954 and MDA-MB-468 cell lines. Primers to the genes of interest were designed to tile across multiple sites in each gene, with several regions enriched for ID4 binding compared to input and IgG controls (Figure 1e and Supplementary Figure 4 for full panel of primers used). ChiP-qPCR validation from independent ChIP experiments validated the findings made from the unbiased ChIP-seq and ChIP-exo approaches. ID4 binding to a number of these loci was also observed in the ID4-positive HCI-002 BLBC patient-derived xenograft (PDX) model (DeRose et al., 2013) (Figure 1f).

To address the specificity of ID4 binding, ChIP-qPCR analysis was conducted on HCC70 cell line modified with the SMARTChoice doxycycline-inducible ID4 short hairpin. Doxycycline treatment resulted in an 87% reduction in ID4 protein expression at 72 h (Supplementary Figure 5). ID4 knockdown reduced ID4-chromatin binding at a majority of sites (Supplementary Figure 6), confirming the specific binding of ID4 to these loci. These findings challenge the proposed classical mechanism of action of ID4 and demonstrates novel ID4-chromatin interactions in BLBC.

Although ID proteins are classically thought to be transcriptional regulators, to date, no studies have reported a systematic analysis of ID4 transcriptional targets. Hence, to determine whether ID4 affects transcription in BLBC, we analysed changes in gene expression using RNA-sequencing following ID4 depletion using the SMARTChoice model described above. This analysis was also used to identify whether ID4 regulates the expression of genes adjacent to ID4 chromatin interactions. Remarkably, differential expression analysis using two analysis methods (Voom and EdgeR) revealed limited changes in gene expression following loss of ID4 (Supplementary Table 2), indicating ID4 does not function as a transcriptional regulator in this model.

The more stringent Voom method, identified a total of six genes as differentially expressed. These genes form a subset of the 21 genes identified using using an independent differential gene expression algorithm called EdgeR. As such, two independent tools reveal almost no ID4 regulated genes. Loss of ID4 did not affect the expression of the ID4-bound genes identified through ChIP-seq and ChIP-exo analysis, as determined through RNA-seq and qPCR (Figure 1g). This suggests that ID4 has a transcription-independent role on chromatin.

### ID4 interacts with DNA damage response proteins

To understand how ID4 interacts with chromatin in the absence of a known DNA interaction domain, endogenous ID4 was purified from asynchronous cells using the proteomic technique Rapid immunoprecipitation Mass spectrometry of Endogenous proteins (RIME) (Mohammed et al., 2013). Anti-ID4 and control IgG were used for immunoprecipitations in biological triplicate (each in technical duplicate or triplicate) followed by mass spectrometry in the HCC70, MDA-MB-468, HCC1954 and OVKATE cell lines. Proteins were considered bone fide ID4-binding targets if they appeared in more than one technical and biological replicate and were not present in any IgG control RIME samples (Figure 2a).

**Figure 2:**
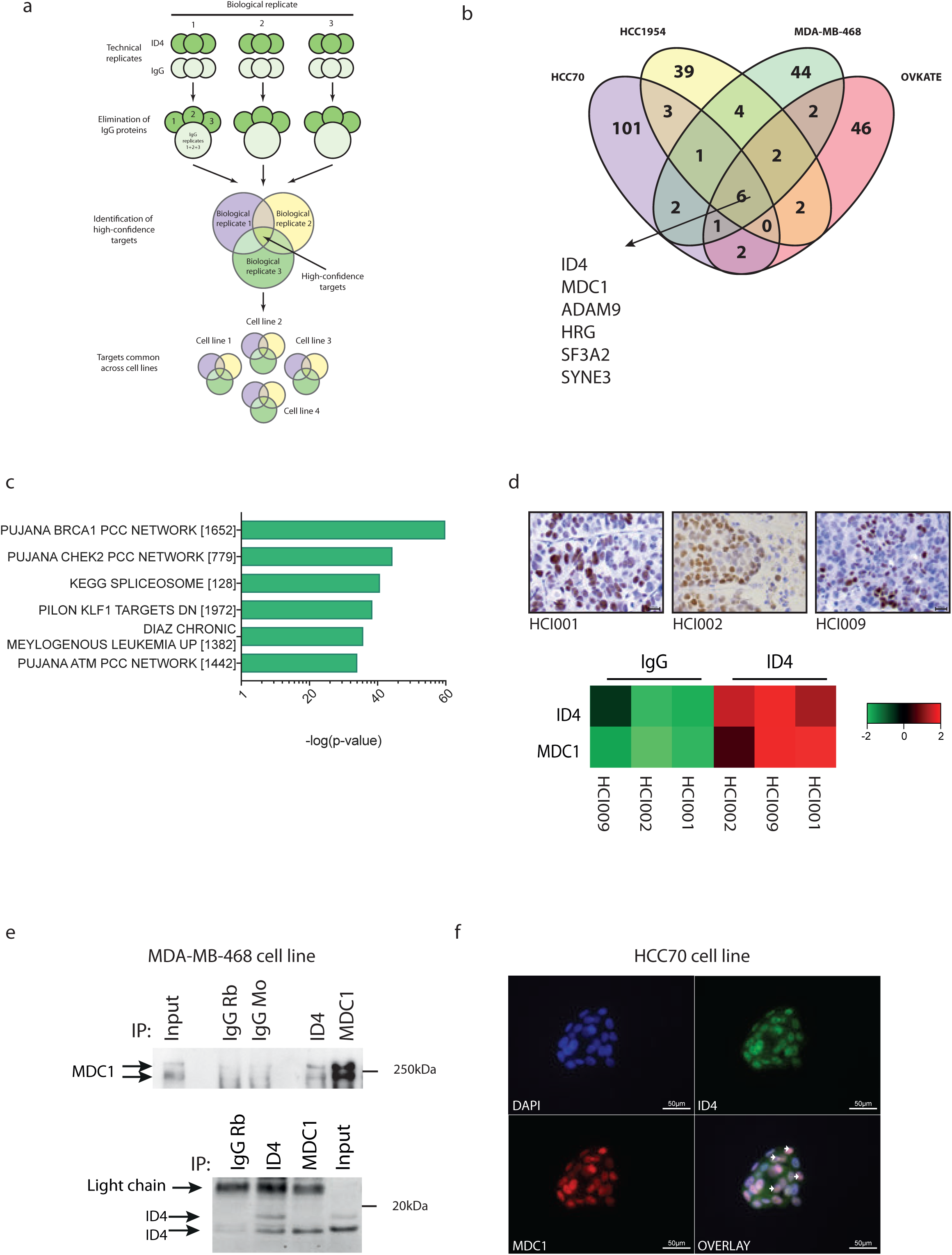
ID4 binds to MDC1 and interacts with the BRCA1 network. A; Schematic of ID4 and IgG RIME data analysis; ID4 and IgG immunoprecipitations were conducted in technical duplicate or triplicate, and in biological triplicate. All proteins identified in the IgG controls (for each biological replicate) were removed from each of the ID4 IPs as non-specific, background proteins. The remaining proteins were compared across technical replicates to generate a list of medium-confidence proteins that were robustly identified in >1ID4 RIME technical replicate. The biological replicates were then compared to generate a list of targets present in >1 biological replicate in an individual cell line (i.e >6 technical replicates conducted over three biological replicates). This resulted in the identification of 22 (HCC70), 21 (HCC1954), 21 (MDA-MB-468) and 30 (OVKATE) proteins. Targets from the different cell lines were compared, identifying a list of ID4 interactors present across multiple cell lines. B; Venn diagram showing comparison of the high-confidence targets identified in all four cell lines; HCC70, HCC1954, MDA-MB-468 and OVKATE. Common targets across the four cell lines are indicated. Overlap generated using Venny (Oliveros, 2007). C; Top six highest enriched gene sets identified through Gene Set Enrichment Analysis (GSEA) of the ID4 proteome (1,106 proteins) identified in (B) (all proteins identified in ID4 RIME and not in IgG RIME), compared to C2 (curated gene sets), C4 (computational gene sets), C6 (oncogenic signatures), C7 (immunologic signatures) and H (hallmark gene sets) gene sets. Proteins identified in >1 technical replicate of ID4 RIME (and not in IgG RIME) were considered in GSEA analysis. D, top; Immunohistochemistry analysis of ID4 protein expression across HCI001, HCI002 and HCI009 triple negative PDX models, 200x magnification, bottom; SWATH proteomic analysis of ID4 and IgG RIME conducted on HCI001, HCI002, HCI009 PDX models. Heatmap showing quantification of ID4 and MDC1 abundance, both significantly differentially expressed proteins in ID4 RIME compared with control IgG RIME (p-value < 0.05 and fold change > 2). RIME and SWATH were conducted on the same sample; one biological replicate per tumour. E; immunoprecipitation was conducted on MDA-MB-468 cells prepared using the RIME protocol. Top: Input control and IgG (mouse and rabbit) compared to IP with anti-ID4 antibodies and MDC1. Western blotting is shown using an independent monoclonal MDC1 antibody. Bottom: Input control and IgG rabbit compared to IP with anti-ID4 antibodies and MDC1. Western blotting is shown using an independent monoclonal ID4 antibody. F; Immunofluorescence analysis of DAPI (blue), ID4 (green) and MDC1 (red) in HCC70 cell line. Example regions of co-localisation indicated with arrows. Scale on images.

RIME analysis identified cell line-unique and common ID4-binding candidates (Figure 2b, Supplementary Table 3). In total 1,106 unique proteins were identified across the four cell lines. Gene Set Enrichment Analysis (GSEA) against the C2 database (4,738 curated gene sets) of the putative ID4 binding partners showed enrichment for chromatin (DNA damage and transcription proteins and histones) and RNA-associated gene sets (including RNA splicing and processing factors), as well as structural and cell cycle related proteins (Figure 2c). This confirms the above ChIP-Seq data showing ID4 interacts with chromatin and nuclear machinery, and may be involved in nuclear processing. The ID4 purified proteome most significantly enriched, resembled the BRCA1-PCC network (PCC: Pearson Correlation Coefficient, 101/852 ID4-associated genes were present in the set of 1652 *BRCA1*-associated genes; p=6.47E-59) (Figure 2c). This gene set comprises networks controlling breast cancer susceptibility (Pujana et al., 2007). Other gene sets originating from this previous study (Pujana et al., 2007), including the CHEK2-PCC and ATM-PCC, were also identified as highly enriched, suggesting an association between ID4 and DNA damage repair factors. We observed only one instance of identification of a bHLH protein, HEB (also known as TCF12). Only one HEB peptide was identified in one of nine technical replicates in the OVKATE cell line. We therefore concluded that HEB was not the primary ID4 interactor, and that ID4 must be acting through novel, non-bHLH mediated mechanisms.

Six proteins were commonly identified in all four-cell lines: ID4 (validating the RIME approach), mediator of DNA damage checkpoint 1 (MDC1), disintegrin and metalloprotease domain 9 protein (ADAM9), Histidine Rich Glycoprotein (HRG), splicing factor 3a subunit 2 (SF3A2) and spectrin repeat containing nuclear envelope family member 3 (SYNE3). The most highly enriched protein, MDC1, was the only protein to be identified in all RIME technical and biological replicates in all four-cell lines. MDC1 has many cellular functions, including cell cycle control and transcription (Townsend et al., 2009, Wilson et al., 2011), yet is best characterised for its role in DNA damage repair, where it associates with DNA (Stewart et al., 2003). Of the other proteins identified, ADAM9, and HRG affect cell motility, SYNE3 is important in cytoskeletal and nuclear organisation (Wilhelmsen et al., 2005) and SF3A2 is required for pre-mRNA splicing and the generation of alternative transcripts (Takenaka et al., 2004). ID4 was previously suggested to bind to mutant p53 and SRSF1 (Fontemaggi et al., 2009, Pruszko et al., 2017), yet no evidence was found of these associations in our analysis, even when one of the same cell lines was evaluated (MDA-MB-468).

To validate these findings in a more clinically relevant setting, we conducted ID4 RIME in PDX models of triple negative breast cancer (DeRose et al., 2013). Three ID4-expressing models (HCI001, HCI002, HCI009) were selected for RIME analysis based on detectable expression of ID4 (Figure 2d). ID4 and MDC1 were the only targets identified in ID4 RIME on all three xenograft models and absent in the IgG RIME (see Supplementary Table 3 for full list of proteins identified in PDX models). This discovery proteomics was supported by quantitative mass spectrometry analysis of the models using the sensitive label-free quantitative proteomic method termed SWATH (Sequential Window Acquisition of all THeoretical mass spectra) (Liebler and Zimmerman, 2013). This analysis showed a high abundance of both ID4 and MDC1 in all three PDX models (Figure 2d). Due to the robust identification of MDC1 in the four cell lines and the subsequent validation of this interaction in three PDX models, we focused further studies on the functional importance of the ID4-MDC1 relationship.

ID4-MDC1 binding was validated using independent methods including immunoprecipitation with an anti-ID4 antibody and anti-MDC1 antibody, followed by Western blot with an independent monoclonal ID4 or a MDC1 antibody (Figure 2e). This Co-IP experiment confirmed the ID4-MDC1 interaction in MDA-MB-468 cells. In addition, immunofluorescence analysis demonstrated co-localisation of ID4 and MDC1 in cell nuclei (Figure 2f and Supplementary Figure 7).

### ID4 interacts with DNA repair proteins at fragile chromatin sites in a DNA damage-dependent manner and regulates DNA damage signaling

To further explore the interaction between ID4 and MDC1 we used proximity ligation assays (PLA), a quantitative measure of protein interaction *in situ*, and confirmed binding of ID4 with MDC1 in MDA-MB-468 cell line (Figure 3a). Imaging reveals that this interaction occurred predominantly in the cell nucleus. Depletion of MDC1 using siRNA resulted in reduction of the PLA signal to control levels. PLA analysis was also used to test the association of ID4 with γH2AX, as it is required for signalling doublestranded DNA breaks and with BRCA1, as it is a critical mediator of the homologous recombination repair pathway (HR). PLA analysis identified enrichment of ID4 binding with MDC1, γH2AX and BRCA1 above control levels in HCC70 and with MDC1 in OVKATE cell lines (Supplementary Figure 8).

**Figure 3:**
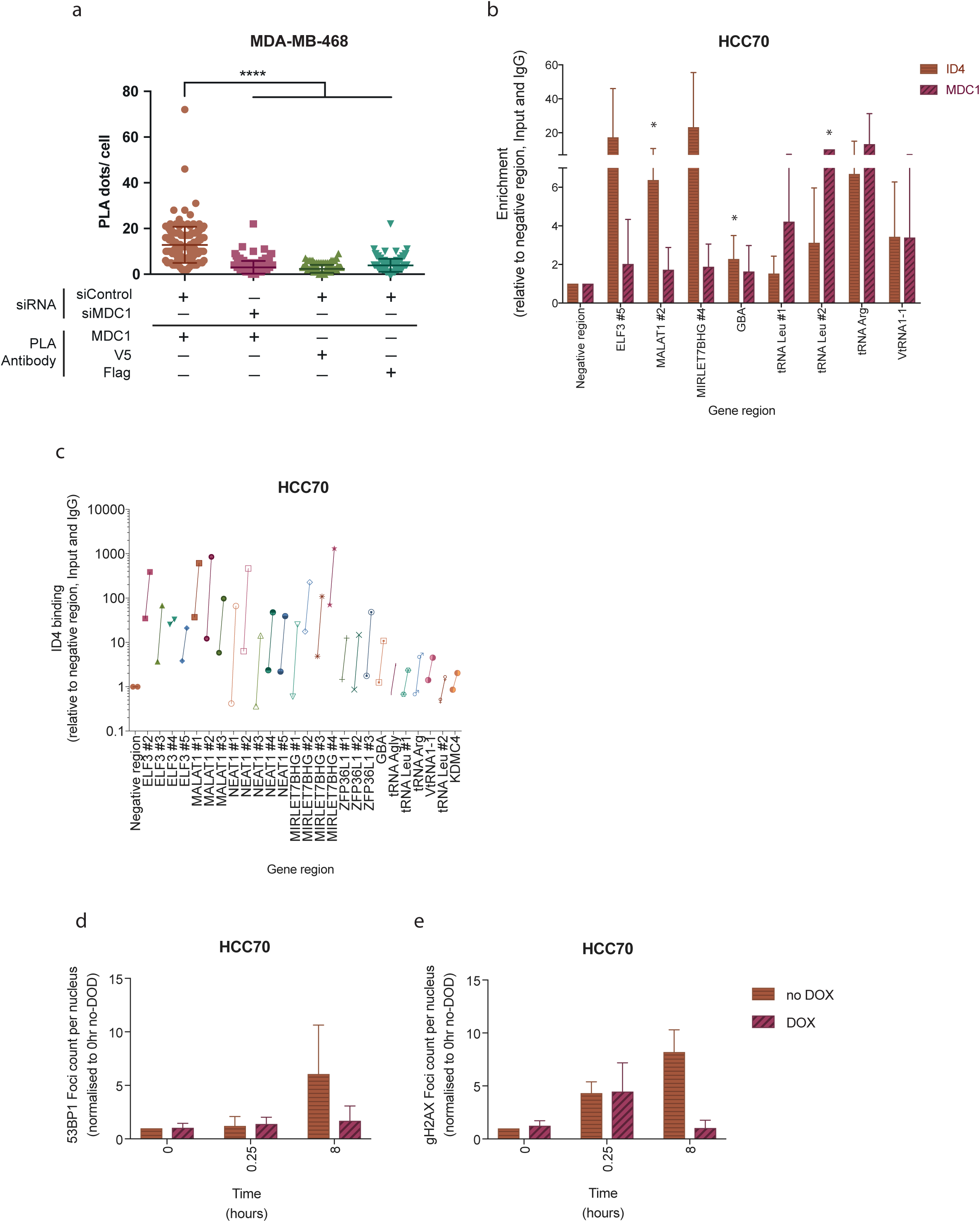
ID4 interacts with DNA damage proteins at fragile chromatin sites in a DNA damage-dependent manner. A; Proximity ligation assay (PLA) analysis of ID4 association with MDC1 in MDA-MB-468 cells. siRNAs targeting MDC1 (siMDCl) and a scrambled control (siControl) used to show specificity of the assay. Graph showing quantification of PLA interactions between ID4 and MDC1 in cells treated with siControl and siMDCl. Data shown in comparison to the interaction between ID4/ V5 and ID4/ FLAG treated with siControl. 50-100 cells captured per condition. Quantification of interactions (number of dots/ cell). B; ChIP-qPCR analysis of ID4 and MDC1 in HCC70 cell line. Binding normalised to input DNA and to a region not bound by ID4 or MDC1 (negative region) and represented as fold-change over IgG control. n=2-5, * indicates p<0.05. C; ChIP-qPCR analysis of ID4 binding in untreated HCC70 cells compared to cells treated with ionising radiation (5Gy with 5 h recovery time prior to fixation). ID4 binding normalised to negative control region, input DNA and IgG control. Data for each gene region is shown for both untreated (left) and ionising radiation-treated (right) cells, connected by a line. Representative of two independent experiments. Quantification of a time course of D: γH2AX and E: 53BP1 DNA damage foci formation following ionising radiation. ID4 was depleted from cells using the SMARTChoice inducible shRNA system following treatment with doxycycline for 72 h prior to treatment with 5 Gy ionising radiation at time 0 and allowed to recover for 0.25 and 8 hours prior to analysis. Four to five images were taken for each condition; 100-200 cells. Number of foci per cell nucleus was calculated using FIJI by ImageJ (Schindelin et al., 2012) and samples were then collated and analysed using the Pandas package in Python 3.5 and graphed using Prism v6. Data is normalised to 0 h, no DOX, no IR time point. n= 3-5.

As ID4 lacks a DNA-binding domain, it is likely that ID4 associates with chromatin through its protein-protein interactions. ChIP-qPCR using antibodies targeting MDC1 demonstrated that the protein co-localises to some of the ID4-bound regions in nonirradiated HCC70 cell line (Figure 3b). By mining published data (Gardini et al., 2014),BRCA1 was also found to bind to chromatin at the regions encoding ZFP36L1, ERRFI1, several tRNAs and the entire NEAT1 and MALAT1 locus in a manner similar to ID4 (Supplementary Figure 2f and g).

To more closely examine the correlation between DNA damage and ID4 complex association with chromatin, we used ionising radiation to induce DNA damage in cells. Irradiation markedly increased ID4 chromatin binding, measured by ChIP-qPCR; up to 160-fold compared to undamaged controls at the NEAT1 gene, and 5-70-fold at ELF3, MALAT1, MIRLET7BHG, ZFP36L1, GBA and various tRNAs (Figure 3c and Supplementary Figure 9). This indicates that ID4 binding is specific and induced by DNA damage. ChIP-qPCR for MDC1 similarly showed induced binding to a majority of these sites in damaged cells (Supplementary Figure 9). This identifies a clear link between 104-chromatin binding and DNA damage.

To investigate the functional role for ID4 in the DDR, DNA damage was induced using ionising radiation and damage sensing was monitored over time. Cells were treated with doxycycline to deplete ID4 for 72h (point of maximal ID4 knockdown) prior to treatment with ionising radiation. Cells were fixed at 0, 15min and 8h following ionising radiation, to examine the DNA damage response over time. As previously reported (Bekker-Jensen et al., 2005), the number of γH2AX foci in control cultures increased at 15min post-IR and remained high at 8h (Figure 3d), while the number of 53BP1 foci increased at 8h following ionising radiation (Figure 3e). Remarkably, upon ID4 depletion, formation of γH2AX and 53BP1 foci was significantly reduced at 8h post-IR. This suggests that ID4 loss impacts DNA damage sensing or repair, revealing a novel role for ID4 in the DNA damage repair response.

### ID4 genetically interacts with *BRCA1* mutation

To determine whether ID4 interacts with DNA repair pathways in clinical disease, we assessed whether there is evidence for genetic mutation of *ID4* in sporadic disease or in the context of familial breast cancer driven by germline *BRCA1* mutation. *ID4* amplification was first evaluated in a cohort of sporadic BLBC. ID4 is reportedly amplified in these cancers (Ren et al., 2012, Network, 2011, Network, 2012), however these results are based on small cohorts and imprecise array Comparative Genomic Hybridisation (aCGH) techniques, both of which affect accurate determination of amplification frequency. To definitively quantify *ID4* amplification frequencies, we used fluorescent *in situ* hybridisation (FISH) in a clinical diagnostic laboratory, a highly specific approach for analysing copy number. FISH was applied to a discovery cohort of 82 oestrogen receptor-negative invasive ductal carcinomas: composed of 42 BLBC (negative for ERα, PR, HER2 and positive for CK5/6, CK14 or EGFR), 14 triple negative non-BLBC (negative for ERα, PR, HER2, CK5/6, CK14 and EGFR) and 26 HER2-Enriched (negative for ERα and PR, positive for HER2) samples (Junankar et al., 2015). *ID4* amplification was exclusive to the BLBC subtype with an amplification frequency of ∼10% (4/42 cases) (Figure 4a). To determine potential genetic interactions between *ID4* and *BRCA1, ID4* copy number and protein expression was evaluated in 42 unselected BLBC (which are not expected to carry *BRCA1* germline mutations) and 97 familial BLBCs from patients with known germline *BRCA1* mutations. *ID4* was amplified at twice the frequency in *BRCA1*-mutant BLBC (∼22%, 21/97 cases) compared to sporadic BLBC (∼10%, 4/42 cases), indicating a selection for *ID4* amplification in cancers arising in patients carrying a mutant *BRCA1* allele (Figure 4c). ID4 protein expression was also significantly higher in the *BRCA1*-mutant BLBC compared to the sporadic BLBC cohort (Figure 4d, Supplementary Figure 10a). *ID4* amplification and protein expression in the *BRCA1*-mutant cohort did not correlate with survival (p = 0.83 and p = 0.89 respectively) (Supplementary Figure 10d and b), likely due to the heterogeneous treatment regimens and small number of amplified cases. High ID4 protein expression in sporadic BLBC has previously been shown to correlate with poor survival (p = 0.0008) (Junankar et al., 2015).

**Figure 4:**
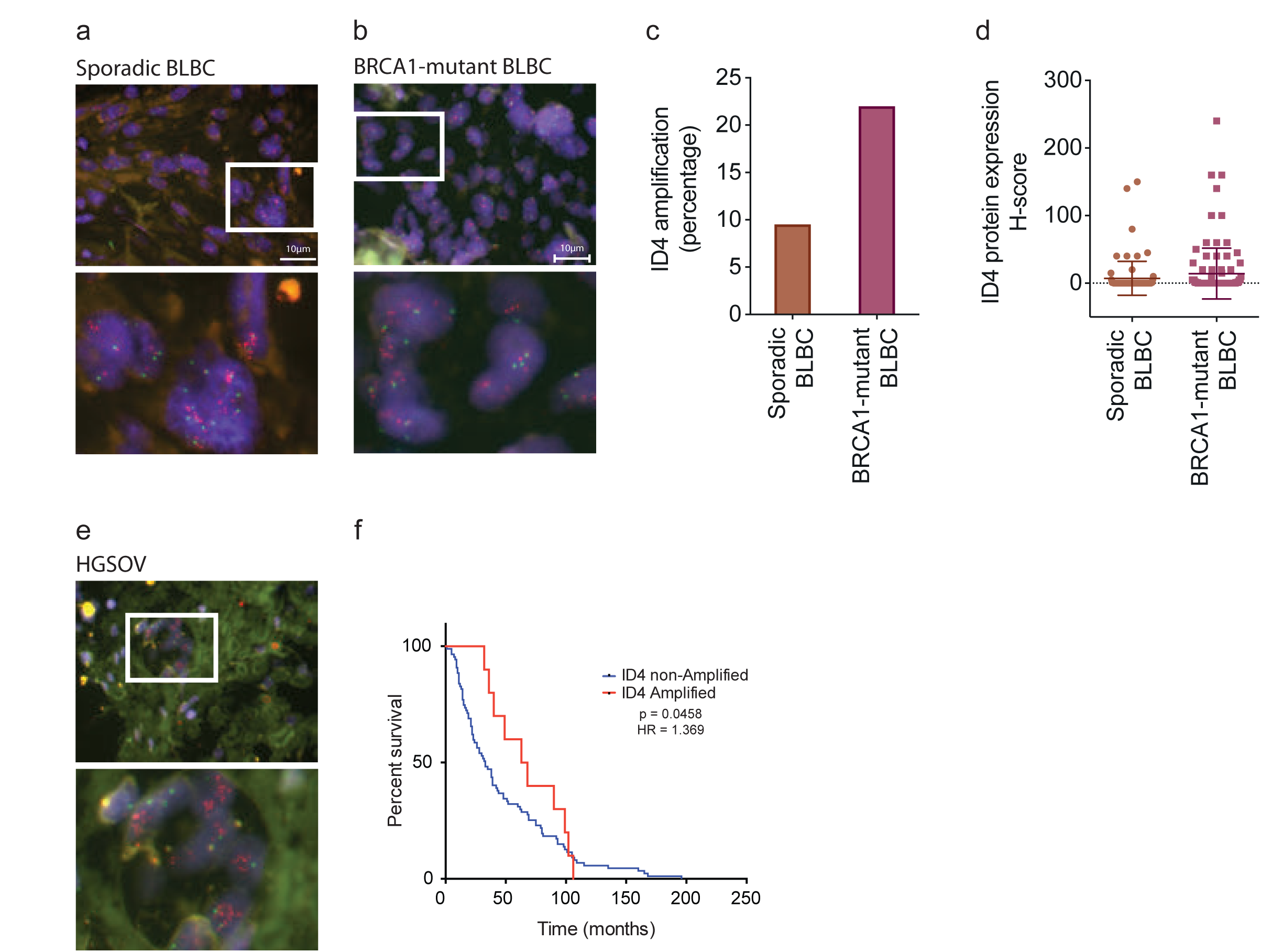
ID4 is selectively amplified in cancers arising in germline BRCA1 mutation carriers. A; ID4 amplification in sporadic and familial BLBC. Example images of fluorescent in situ hybridisation (FISH) analysis of ID4 (red) and CEP6 centromeric control (green) copy number aberrations in a cohort of 42 sporadic BLBCs. B; Example images of FISH analysis of ID4 (red) and CEP6 centromeric control (green) copy number aberrations in a cohort of 97 *BRCA1*-mutant BLBCs. C; Percentage of ID4 amplified cases in sporadic and *BRCA1*-mutant BLBCs presented in (A) and (B). p<0.03 (Two sample t-test). D; Immunohistochemical analysis of ID4 protein expression in the sporadic and *BRCA1*-mutant BLBCs presented in (A) and (B). ID4 protein quantified using the H-score method. H-score = intensity of staining (on a scale from 0-3) x percentage of cells positive. p=0.007 (Mann-Whitney two-tailed test). E; Example images of FISH analysis of ID4 (red) and CEP6 centromeric control (green) copy number aberrations in a cohort of 97 high-grade serous ovarian cancers. F; Kaplan-Meier survival analysis of HGSOV patients grouped according to amplification status, p = 0.046, HR = 1.369.

Our biochemical and molecular cell line data suggests that ID4 has similar function in BLBC and HGSOV, so we examined whether ID4 is also amplified in HGSOV. Analysis of 97 sporadic HGSOV cases (Montavon et al., 2012) showed a comparable rate of *ID4* amplification relative to sporadic BLBC; ∼10% (10/97 in cases of HGSOV) (Figure 4e, Supplementary Figure 10). However, these frequencies are lower than that published using Array CGH (Network, 2011, Network, 2012), potentially suggesting that aCGH methods overestimate amplification frequencies. *ID4* amplification in HGSOV correlated significantly with longer overall survival (65.5 months) compared to nonamplified patients (33.0 months) (p = 0.046, HR = 1.37, 95% CI = 0.76 to 2.46) (Figure 4f). This analysis corresponds with published data in The Cancer Genome Atlas, although there it fails to reach statistical significance (58.1 and 44.1 month survival, respectively, p = 0.07) (Network, 2011) (Supplementary Figure 11e). ID4 protein expression correlates with amplification (r = 0.457 and p <0.0001) (Supplementary Figure 11d), though did not correlate with patient survival in the HGSOV cohort (Supplementary Figure 11c, p = 0.225). This finding, in a cohort treated primarily with platinum, may suggest an association between ID4 amplification and sensitivity to platinum therapy (Montavon et al., 2012).

## Discussion

By elucidating the genetic drivers and dependencies of BLBC, we aim to improve our understanding of this complex, heterogeneous disease, potentially leading to the identification of novel targets and therapeutics. ID4 is required for normal mammary gland development (Junankar et al., 2015, Best et al., 2014, Dong et al., 2011) and acts as a proto-oncogene in BLBC (Beger et al., 2001, Branham et al., 2016, Crippa et al., 2014, de Candia et al., 2006, Junankar et al., 2015, Roldán et al., 2006, Shan et al., 2003, Thike et al., 2015, Wen et al., 2012). Depletion of ID4 reduces BLBC cell line growth *in vitro* and *in vivo* (Junankar et al., 2015), suggesting that ID4 controls critical, yet unknown pathways in BLBC.

We have conducted the first systematic mapping of the chromatin interactome of ID4, or indeed any ID protein, in mammalian cells. Using ChIP-seq we have identified novel, reproducible ID4 binding sites within the BLBC genome. ID4 bound to large regions of chromatin, up to 10kb in length, at a very small number of loci, suggesting that ID4 is not binding as part of a transcriptional regulatory complex. This conclusion is supported by the observation that ID4 knockdown did not affect gene expression at these loci. The regions identified through ChIP-seq and ChIP-exo were typically the bodies of genes that are highly transcribed and mutated in cancer (Seo et al., 2012, Natale et al., 2013, Meier et al., 2007), characteristics of fragile, DNA damage-prone sites. Interestingly, these sites primarily encode non-coding RNA, including lncRNA, microRNA precursors and tRNA. The lncRNA NEAT1 and MALAT1 are some of the most abundant cellular RNAs and the genes encoding them undergo recurrent mutation in breast cancer (Nik-Zainal et al., 2016). Furthermore, they are upregulated in ovarian tumour cells and are associated with higher tumour grade and stage and in metastases (Vafaee et al., 2017). The genomic loci encoding tRNAs are also highly transcriptionally active, co-localising with DNaseI hypersensitivity clusters and transcription factor binding sites, including BRCA1 and POLR2A (Gardini et al., 2014). Transcriptional activity is highly stressful and associated with genomic stress and DNA damage (Seo et al., 2012). Together these data suggest that ID4 binds preferentially to sites of active transcription and DNA damage.

Systematic and unbiased RIME proteomics revealed hundreds of ID4 interacting proteins, which were highly enriched for *BRCA1*-associated proteins. Interestingly, no bHLH proteins were reproducibly found in complex with ID4, although they are the canonical ID binding partners in certain non-transformed cells (Benezra et al., 1990, Jen et al., 1992, Loveys et al., 1996, Langlands et al., 1997, Roberts et al., 2001). This is unlikely to be a technical artefact, as ID4-bHLH interactions are readily detected in non-transformed mammary epithelial cells using the same method (H. Holliday; Unpublished data). Rather, ID4 may have alternate binding partners as a consequence of downregulation of bHLH proteins E2A, HEB and ITF2 in BLBC (H. Holliday; Unpublished data). ID4 has previously been shown to interact with mutant p53 and SRSF1 (Fontemaggi et al., 2009, Pruszko et al., 2017), yet these two proteins were not identified through RIME analysis across four breast and ovarian cancer cell lines, including MDA-MB-468 cell line, which was previously used to identify ID4-mutant p53 and ID4-SRSF1 interactions (Fontemaggi et al., 2009). This disparity may be the consequence of methodological differences.

Five novel proteins were found to interact with ID4 with high confidence in all 4 models examined, namely MDC1, ADAM9, HRG, SF3A2 and SYNE3. These warrant further investigation to clarify the role they may have in the mechanism of action of ID4.

MDC1, the most reproducible binding partner of ID4 identified through RIME analysis, forms a quasi HLH domain (Stucki et al., 2005), which may enable interaction with the HLH domain of ID4. MDC1 is recruited to sites of DNA damage to amplify the phosphorylation of H2AX (to form γH2AX) and recruit downstream signalling proteins (Lukas et al., 2004, Stewart et al., 2003). Deficiency in MDC1, much like BRCA1, results in hypersensitivity to double-stranded DNA breaks (Ando et al., 2013, Lou et al., 2006). MDC1 was also recently found to associate with ID3, suggesting that this is a conserved feature of ID proteins (Lee et al., 2017). MDC1 also interacted with many of the sites of chromatin interaction by ID4, and this was increased following DNA damage. Using the sensitive PLA method, we also find ID4 in close proximity of known MDC1 interactors, including BRCA1 and gamma-H2AX. These data suggest a model in which ID4 associates with the DNA damage repair apparatus at sites of genome instability or damage via its interaction with MDC1. Importantly, this association affects the DNA damage response as the number of HR (BRCA1) and NHEJ (p53BP1) foci are reduced following ID4 depletion (Liu et al., 2012, Lou et al., 2006, Lukas et al., 2004, Stewart et al., 2003, Stucki et al., 2005, Zhang et al., 2012, Lu et al., 2013). As complex feedback mechanisms govern the DNA damage response, further investigation is required to determine whether ID4 association with MDC1 ultimately promotes or impedes DNA repair.

These data suggest a remarkable gain of function by ID4 in cancer, switching from a role in transcription in normal mammary epithelial cells (Best et al., 2014, Junankar et al., 2015) to a role in DNA damage in BLBC. A similar dichotomy of function has been described for BRCA1, which has proposed functions as a transcription factor in nontransformed cells (Furuta et al., 2005, Gardini et al., 2014), but a primary role in DNA repair in cancer (Scully et al., 1997, Cortez et al., 1999). Transcription is a stressful cellular process causing significant DNA damage and repair (Seo et al., 2012, Malewicz and Perlmann, 2014). Thus, a role for transcription factors in localising DNA damage machinery to chromatin may be an important cellular capability.

ID4 and BRCA1 have been shown to inversely correlate in expression (de Candia et al., 2006, Roldán et al., 2006, Wen et al., 2012, Crippa et al., 2014, Junankar et al., 2015, Branham et al., 2016): however, this is the first demonstration of a biochemical interaction of ID4 with DNA damage foci and the DNA damage repair apparatus.

Mutations in BRCA1 predispose carriers predominantly to cancers of the breast and ovaries (mostly BLBC and HGSOC), though the mechanism driving tumorigenesis in these patients is still unclear. In addition to the biochemical interaction between ID4 and BRCA1, we also show a genetic interaction, in that *ID4* is amplified at twice the frequency in *BRCA1*-mutant BLBC compared to sporadic BLBC.

A defect in BRCA1 function in the absence of germline BRCA1 mutations, termed BRCAness (Konstantinopoulos et al., 2010, Lips et al., 2013, Liu et al., 2014, Turner et al., 2004) correlates with sensitivity to platinum-based chemotherapies and PARP inhibitors, both traditionally used to treat *BRCA1*-mutant cancer, especially HGSOC.Consistent with this proposal, ID4 expression correlates with BRCAness in breast cancer (Branham et al., 2016) and we now show that women with /D4-amplified HGSOC, treated primarily with platinum-based agents, have improved survival. Together, these observations lead us to postulate that overexpression of ID4 may function to suppress HR function, contributing to a BRCA-ness phenotype in BRCA1-wild type cells. Cells with heterozygous *BRCA1* mutation may be further susceptible to this function of ID4, leading to selection for *ID4* amplification in familial cases. Further work is needed to test the hypothesis that ID4 expression may predict, or even regulate, sensitivity to therapies targeting HR deficiency, such as platinum-containing chemotherapy and PARPi.

## Supplementary Figure Legends

## Materials and methods

### Mammalian cell culture growth conditions

Cell lines were obtained from American Type Culture Collection and verified using cell line fingerprinting. HCC70 and HCC1937 cell lines were cultured in RPMI 1640 (Thermo Fisher Scientific) supplemented with 10% (v/v) FBS, 20 mM HEPES (Thermo Fisher Scientific) and 1 mM Sodium Pyruvate (Thermo Fisher Scientific). MDA-MB-468 and OVKATE cell lines were cultured in RPMI 1640 (Thermo Fisher Scientific) supplemented with 10% (v/v) FBS, 20 mM HEPES (Thermo Fisher Scientific) and 0.25% (v/v) insulin (Human) (Clifford Hallam Healthcare). Cells stably-transduced with SMARTChoice lentiviral vectors were grown in the presence of 1 μg/mL puromycin.

### Imaging

Immunohistochemistry images were obtained using an inverted epifluorescence microscope (Carl Zeiss, ICM-405, Oberkochem, Germany). Images were captured by the Leica DFC280 digital camera system (Leica Microsystems, Wetzlar, Germany). The Leica DM 5500 Microscope with monochrome camera (DFC310Fx) or Leica DMI SP8 Confocal with 4 lasers (405 nm, 488 nm, 552 nm and 638 nm) and two PMT detectors were used to capture standard fluorescent and confocal images.

### SMARTChoice inducible lentiviral system

ID4 and control lentiviral shRNA constructs (SMARTchoice) were purchased commercially (Dharmacon, GE, Lafayette, CO, USA). Successfully transfected cells were selected using puromycin resistance (constitutive under the humanEF1a promoter). For ID4 knockdown analysis, HCC70 cells with SMARTChoice shID4 #1 #178657 (VSH6380-220912204), SMARTChoice shID4 #2 #703009 (VSH6380-221436556), SMARTChoice shID4 #3 #703033 (VSH6380-221436580), SMARTChoice shNon-targeting (VSC6572) and mock-infected cells. Cells were treated with vehicle control or with 1 μg/mL doxycycline for 72 hours before harvesting protein and RNA directly from adherent cells. The SMARTChoice shID4 #2 #703009 (VSH6380-221436556) produced the highest level of ID4 knockdown and was used for further analysis.

### Non-lethal DNA damage induction with ionising radiation

Cells were seeded at 2.5 ×10^5^ (HCC70) or 2.2 ×10^5^ (MDA-MB-468) cells/well in a 6-well plate in normal growth medium. One day post seeding, cells were exposed to 2-5 Gy of ionising radiation using an X-RAD 320 Series Biological Irradiator (Precision X-Ray, CT, USA). Cells were returned to normal tissue culture incubation conditions and harvested at designated time points.

### Gene expression analysis

Total RNA was prepared for using the miRNeasy RNA extraction kit (Qiagen), according to the manufacturer′s instructions. cDNA was generated from 1000ng RNA using the Transcriptor First Strand cDNA Synthesis Kit (Roche) using oligo-dT primers and following the manufacturer’s instructions. qPCR analysis was used to analyse mRNA expression levels using Taqman probes (Applied Biosystems/Life Technologies) as per manufactures specifications (Table 2) using an ABI PRISM 7900 HT machine. qPCR data was analysed using the Δ ΔCt method (Schefe et al., 2006).

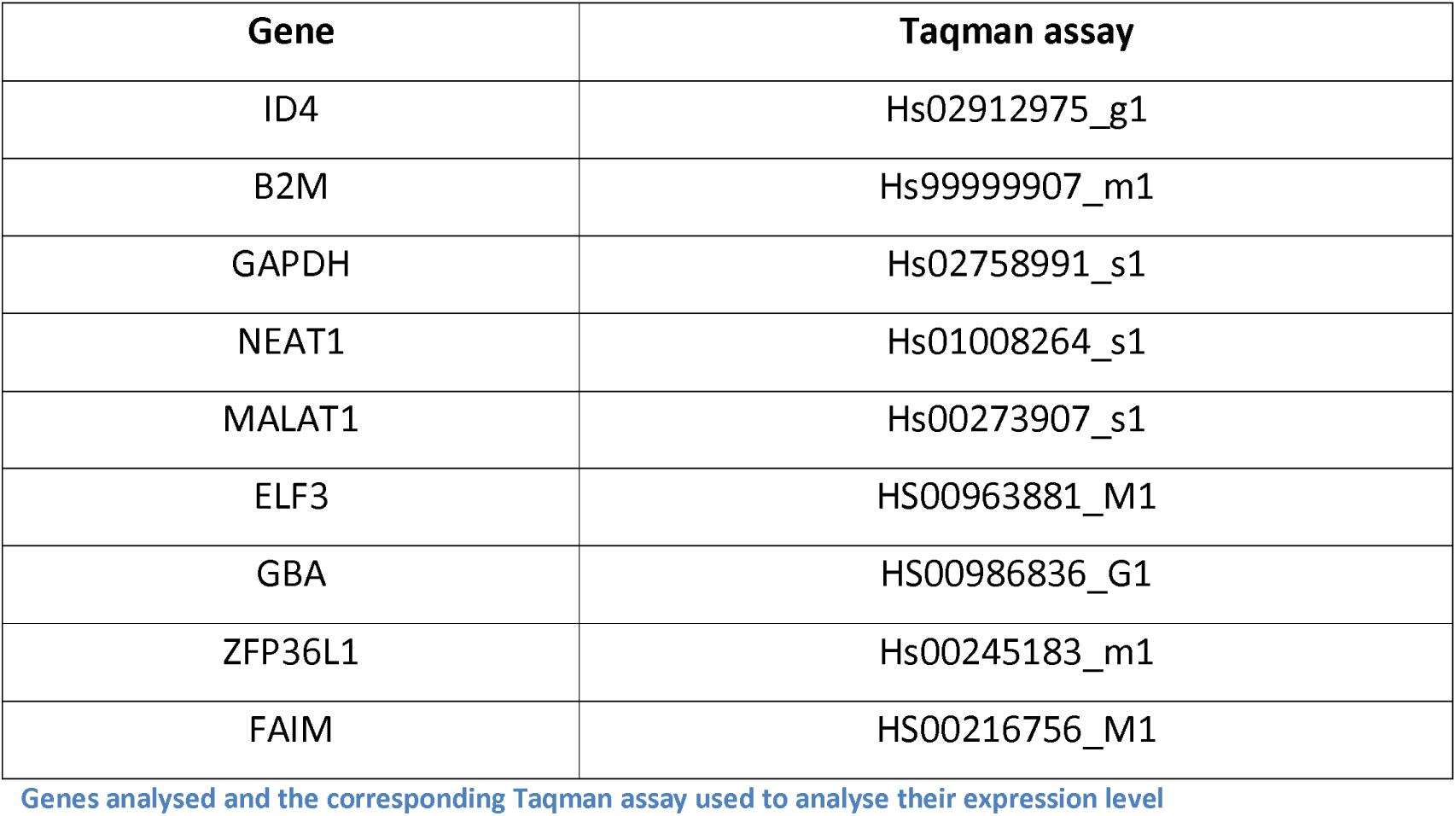

### Genes analysed and the corresponding Taqman assay used to analyse their expression level

#### Protein analysis

Cells were lysed, unless specified, using RIPA [0.88% (w/v) Sodium Chloride, 1% (v/v) Triton X-100, 0.5% (w/v) Sodium Deoxycholate, 0.1% (w/v) SDS, 0.61% (v/v) Tris (Hydroxymethyl)Aminomethane and protease and phosphatase inhibitors (Roche)] or Lysis Buffer 5 (10 mM Tris pH 7.4,1 mM EDTA, 150 mM NaCl, 1% Triton X-100 and protease and phosphatase inhibitors). If required, protein was quantified using the Pierce BCA Protein Assay Kit (Thermo Fisher Scientific) according to manufacturer’s instructions. Western blotting analysis was conducted as previously described (Junankar et al., 2015). MDC1 protein was analysed using 3-8% tris/acetate gels and PVDF nitrocellulose membrane for MDC1 analysis (BioRad). All other proteins were analysed using the LiCor Odyssey system (Millenium Science, Mulgrave, VIC, Australia). Protein expression was analysed using antibodies targeting ID4 (Biocheck anti-ID4 rabbit monoclonal BCH-9/82-12, 1:40,000), [β-Actin(Sigma anti-Actin mouse monoclonal A5441, 1:5,000) and MDC1 (Sigma anti-MDC1 mouse monoclonal M2444, 1:1000).

#### Co-immunoprecipitation

Co-immunoprecipitations (IP) were conducted using 10 [μL per IP Pierce Protein A/G magnetic beads (Thermo Fisher Scientific) with 2 [μg of antibody: IgG rabbit polyclonal (Santa Cruz sc-2027), ID4 (1:1 mix of rabbit polyclonal antibodies: Santa Cruz L-20: sc-491 and Santa Cruz H-70: sc-13047), MDC1 (rabbit polyclonal antibody Merck Millipore #ABC155). Beads and antibody were incubated at 4°C on a rotating platform for a minimum of 4 h. Beads were then washed three times in lysis buffer before cell lysate was added to the tube. Lysates were incubated with beads overnight at 4°C on a rotating platform. Beads were washed three times in lysis buffer and resuspended in 2 x NuPage sample reducing buffer (Life Technologies) and 2 × NuPage sample running buffer (Life Technologies) and heated to 85°C for 10 min. Beads were separated on a magnetic rack and supernatant was analysed by western blotting as described above.

#### Rapid Immunoprecipitation and Mass Spectrometry of Endogenous Proteins (RIME)

Cells were fixed using paraformaldehyde (PFA) (ProSciTech, Townsville, QLD, Australia) and prepared for RIME (Mohammed et al., 2013) and ChIP-seq (Schmidt et al., 2009, Serandour et al., 2013) as previously described. Cross-linking was performed for 7 min for RIME experiments and 10 min for ChIP-seq, ChIP-exo and ChIP-qPCR experiments. Samples were sonicated using a Bioruptor Standard (Diagenode, Denville, NJ, USA) for 30-35 cycles of 30 sec QN/30 sec OFF (sonication equipment kindly provided by Prof. Merlin Crossley, UNSW). IP was conducted on 60 (ChIP-seq/ ChIP-exo) to 120 (RIME) million cells using 100 μL beads/20 μg antibody. Correct DNA fragment size of 100-500bp was determined using 2% agarose gel electrophoresis.

Patient-derived xenograft tumour models were cross-linked at 4°C for 20 min in a solution of 1% Formaldehyde (ProSciTech), 50 mM Hepes-KOH, 100 mM NaCl, 1 mM EDTA, 0.5 mM EGTA and protease inhibitors (H. Mohammed, personal communication). Samples (0.5 mg of starting tumour weight) were dissociated using a Polytron PT 1200E tissue homogeniser (VWR) and sonicated using the Branson Digital Sonifer probe sonicator (Branson Ultrasonics, Danbury, CT, USA) with a microtip attachment for 3-4 cycles of 10 x [0.1 sec ON, 0.9 sec OFF]

**Mass spectrometry analysis** was conducted at the Australian Proteomic Analysis Facility (APAF) at Macquarie University (NSW, Australia (Huang et al., 2015). Briefly, samples were denatured in 100 mM triethylammonium bicarbonate and 1% w/v sodium deoxycholate, disulfide bonds were reduced in 10 mM dithiotreitol, alkylated in 20 mM iodo acetamide, and proteins digested on the dynabeads using trypsin. After C18 reversed phase (RP) StageTip sample clean up, peptides were submitted to nano liquid chromatography coupled mass spectrometry (MS) (nanoLC-MS/MS) characterisation. MS was performed using a TripleToF 6600 (SCIEX, MA, USA) coupled to a nanoLC Ultra 2D HPLC with cHipLC system (SCIEX). Peptides were separated using a 15 cm chip column (ChromXP C18, 3 μm, 120 Å)(SCIEX). The mass spectrometer was operated in positive ion mode using a data dependent acquisition method (DDA) and data independent acquisition mode (DIA or SWATH) both using a 60 min acetonitrile gradient from 5 - 35%. DDA was performed of the top 20 most intense precursors with charge stages from 2+ - 4+ with a dynamic exclusion of 30 s. SWATH-MS was acquired using 100 variable precursor windows based on the precursor density distribution in data dependent mode. MS data files were processed using ProteinPilot v.5.0 (SCIEX) to generate mascot generic files. Processed files were searched against the reviewed human SwissProt reference database using the Mascot (Matrix Science, MA, USA) search engine version 2.4.0. Searches were conducted with tryptic specificity, carbamidomethylation of cysteine residues as static modification and the oxidation of methionine residues as a dynamic modification. Using a reversed decoy database, false discovery rate was set to less than 1% and above the Mascot specific peptide identity threshold. For SWATH-MS processing, ProteinPilot search outputs from DDA runs were used to generate a spectral library for targeted information extraction from SWATH-MS data files using PeakView v2.1 with SWATH MicroApp v2.0 (SCIEX). Protein areas, summed chromatographic area under the curve of peptides with extraction FDR ≤ 1%, were calculated and used to compare protein abundances between bait and control IPs.

### Chromatin immunoprecipitation-quantitative real-time PCR analysis

Chromatin immunoprecipitation (ChIP) was conducted as described previously (Schmidt et al., 2009); however, following overnight IP, the samples were processed using a previously described protocol (Nguyen et al., 2001, Taberlay et al., 2011). DNA was purified then quantified using quantitative real-time PCR analysis. Control regions analysed and primers used are listed in Supplementary Table 4.

Relative enrichment of each region/primer set was calculated by taking an average of each duplicate reaction. The input Ct value was subtracted from the sample Ct value and the Ct converted using the respective PE for each primer set. The relative ChIP enrichment is then calculated by dividing the gene region of interest by the specific control region that is negative for both ID4 and H3K4Me3 binding (IFF01/NOP2 #1 primer). The formula for this normalisation is below:

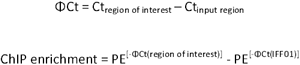

A sample was considered to be enriched if the fold-change over IgG control for each region was >2.

### Chromatin immunoprecipitation-sequencing

**Chromatin immunoprecipitation and sequencing** (ChIP-seq) was conducted as previously described (Schmidt et al., 2009). Samples were prepared and sequenced at Cancer Research United Kingdom (CRUK), Cambridge, UK. Antibody conditions for ChIP are the same as those used for RIME, with the addition of antibodies targeting H3K4Me3 (Active Motif #39159) and γH2AX (Ser139) (1:1 mix of Cell Signalling #2577 and Merck Millipore clone JBW301). Samples were sequenced at CRUK using an Illumina HiSeq 2500 single-end 50 bp sequencing. Quality control was conducted using FastQC (Andrews, 2010) and sequencing adapters trimmed using cutadapt (Martin, 2011).Reads were aligned using Bowtie for lllumina vO.12.7 (Langmead and Salzberg, 2012)followed by Sam-to-Bam conversion tool (Li et al., 2009) and alignment using Bwa v0.705a (Li et al., 2009). Alignment statistics were generated using samtools flagstat [version 0.1.18 (r982:295)] (Li et al., 2009). ChIP-seq peaks were called using the peak calling algorithm HOMER v4.0 and MACS v1.4.2 (Shi et al., 2008, Zhang et al., 2008).

**Chromatin Immunoprecipitation-exonuclease sequencing** (ChIP-exo) was conducted as previously described (Serandour et al., 2013). Samples were prepared and sequenced at CRUK

### qPCR analysis of ChIP DNA

Publicly available H4K3Me3 ChIP-sequencing data and the ID4 ChIP-sequencing data generated in this project were visualised using UCSC Genome Browser (genome.ucsc.edu and (Kent et al., 2002)). Regions of positive and negative enrichment were selected and the 500-1,000 bp DNA sequence was imported into Primer3, a primer design interface, web version 4.0.0 (Koressaar and Remm, 2007, Untergasser et al., 2012). Primers were designed with a minimum primer amplicon length of 70 bp. Primers were confirmed to align with specific DNA segments by conducting an *in silico* PCR using UCSC Genome Browser (genome.ucsc.edu and (Kent et al., 2002)). Oligo primers were ordered from Integrated DNA Technologies (Singapore). Primers were tested to determine adequate primer efficiency (between 1.7-2.3). All assays were set up using an EPmotion 5070 robot (Eppendorf, AG, Germany) and run on an ABI PRISM 7900 HT machine (Life Technologies, Scoresby, VIC, Australia). Briefly, reactions were performed in triplicate in a 384-well plate. Each reaction consisted of 1 [μL 5 [μM Forward primer, 1 [μL 5 [μM Reverse primer, 5 *[μl* SYBR Green PCR Mastermix (Thermo Fisher) and 3 [μL DNA. A standard curve was created using unsonicated, purified DNA extracted from the HCC70 cell line in 10-fold dilutions (1, 0.1, 0.01, 0.001, 0.0001).

PCR cycling was as follows: 1 cycle at 50°C for 2 min, 1 cycle at 95°C for 10 min, followed by 40 cycles of 95°C for 15 sec and 60°C for 1 min. A dissociation step was conducted at 95°C for 15 sec and 60°C for 15 sec. Data was analysed and a standard curve created using SDS 2.3 software (Applied Biosystems). The slope was used to calculate the PE using the qPCR Primer Efficiency Calculator (Thermo Fisher Scientific, available at thermofisher.com).

### Patient-derived xenograft and histology

All experiments involving mice were performed in accordance with the regulations of the Garvan Institute Animal Ethics Committee. NOD.CB17-Prkdc^scid^/Arc mice were sourced from the Australian BioResources Ltd. (Moss Vale, NSW, Australia). Assoc. Prof Alana Welm (Oklahoma Medical Research Foundation) kindly donated the patient-derived xenografts (PDX) models used in this study. Models were maintained as described elsewhere (DeRose et al., 2013). Tumour chunks were transplanted into the 4^th^ mammary gland of 5-week old recipient NOD.CB17-Prkdc^scid^/Arc mice. Tumours were harvested at ethical endpoint, defined as having a tumour approximately 1 mm^3^ in size or deterioration of the body condition score. At harvest, a cross-section sample of the tumour was fixed in 10% neutral buffered formalin (Australian Biostain, Traralgon, VIC, Australia) overnight before transfer to 70% ethanol for storage at 4°C before histopathological analysis. The formalin fixed paraffin embedded (FFPE) blocks were cut in 4 [μm-thick sections and stained for ID4 (Biocheck BCH-9/82-12, 1:1,000 for 60 min following antigen retrieval using pressure cooker 1699 for 1 min, Envision Rabbit secondary for 30 min). Protein expression was scored by a pathologist using the H-score method (Budwit-Novotny et al., 1986).

### Fluorescent In Situ Hybridisation

Tissue sections were analysed using Fluorescent In Situ Hybridisation (FISH) to examine the genomic region encoding *ID4* (6p22.3). *ID4* FISH Probe (Orange 552 nm-576 nm, Empire Genomics, NY, USA) was compared to the control probe *CEP6* (Chromosome 6, Green 5-Fluorescein dUTP). This *CEP6* probe marks a control region on the same chromosome as *ID4* and is used to normalise *ID4* copy number. Breast pathologist Dr Sandra O’Toole oversaw the FISH quantification for all samples.

### Immunofluorescence and Proximity ligation assays

#### Immunofluorescence

Cells were seeded on glass coverslips (Coverglass, 13 mm, VITLAB, Germany). At harvest, media was removed, cells were washed twice with PBS without salts and fixed in 4% paraformaldehyde (PFA) (ProSciTech) for 10 min. Cells were again washed twice with PBS without salts (Thermo Fisher Scientific) before permeabilising for 15 min with 1% Triton-X (Sigma-Aldrich) in PBS and then blocking with 5% BSA in PBS without salts for 1 h at room temperature. Cells were washed twice with PBS without salts and antibodies were applied overnight at 4°C: ID4 (Biocheck BCH-9/82-12, 1:1,000), MDC1 (Sigma-Aldrich M2444, 1:1,000), BRCA1 (Merck Millipore (Ab-1), MS110,1:250), γH2AX (Ser139) (Merck Millipore clone JBW301 05-636,1:300), FLAG (Sigma-Aldrich M2, 1:500) and V5 (Santa Cruz sc-58052, 1:500). Cells were washed twice with PBS without salts then secondary antibodies were applied for 1 h at room temperature. Cells were washed twice with PBS without salts, with the second wash containing DAPI (1:500 dilution) and phalloidin (1:1,000 dilution) (CytoPainter Phalloidin-iFluor 633 Reagent Abcam ab176758). Cells were then mounted on slides using 4 [μL of Prolong Diamond (Thermo Fisher Scientific).

#### Duolink Proximity ligation assay analysis (PLA)

PLA was conducted using Duolink PLA technology with Orange mouse/ rabbit probes (Sigma-Aldrich, DU092102) according to the manufacturers instructions. Images were captured using SP8 6000 confocal imaging with 0.4um Z-stacks. Maximum projects were made for each image (100-200 cells) and quantified using FIJI by ImageJ (Schindelin et al., 2012) as described previously (Law et al., 2017). Quantification was conducted on a minimum of 50 cells. Data us represented as number of interactions (dots) per cell.

#### Quantification of DNA damage foci

Image quantification was conducted using FIJI v2.0.0 image processing software (Fiji is just ImageJ, available at Fiji.sc, (Schindelin et al., 2012)) as previously described (Law et al., 2017). Four to five images were taken of each sample. The DAPI channel was supervised to enable accurate gating of cell nuclei for application to other channels. Size selection (pixel size 2,000 to 15,000) and circularity (0.30-1.00) cut-offs were used. Cells on the edge of the image were excluded from the analysis. The number of DNA damage foci per cell nucleus was calculated for approximately 100-200 cells. The information for individual samples was then collated and analysed using the Pandas package in Python 3.5.

### Clinical Cohorts

#### Basal-like breast cancer

Samples were stratified into groups as follows: 42 BLBC (negative for ERα, PR, HER2 and positive for CK5/6, CK14 or EGFR), 14 triple negative non-BLBC (negative for ERα, PR, HER2, CK5/6, CK14 and EGFR) and 26 HER2-Enriched (negative for ERα and PR, positive for HER2). *BRCA1*-mutation status in this cohort us unknown, however it is expected to occur in approximately 6.5% of BLBC patients (Hartman et al., 2012). Samples were obtained under the Garvan Institute ethical approval number HREC 08/145.

#### Kathleen Cuningham Foundation Consortium for research into familial breast cancer (KConfab)

*BRCA1*-mutant BLBC was sourced from KConfab. A total of 97 BRCA1-mutant BLBC cases were obtained under the Garvan Institute ethical approval number HREC 08/145.

#### Ovarian Cancer

A total of 97 HGSOV cases were obtained under the Human Research Ethics Committee of the Sydney South East Area Hospital Service Northern Section (00/115) (Montavon et al., 2012).

## Acknowledgements

We thankfully acknowledge the support of the following funding bodies: Australia Postgraduate Award (to LAB and HH), Boehringer Ingelheim Fonds, Cancer Council NSW, The National Health and Medical Research Council of Australia, The McMurtrie family, Beth Yarrow Memorial Award, Castle Harlan Award, LH Ainsworth Award and Grant Houghton for assistance with data analysis. Research conducted at APAF was supported by the Australian Government’s National Collaborative Research Infrastructure Scheme.

---

**Supplementary Figure 1:**
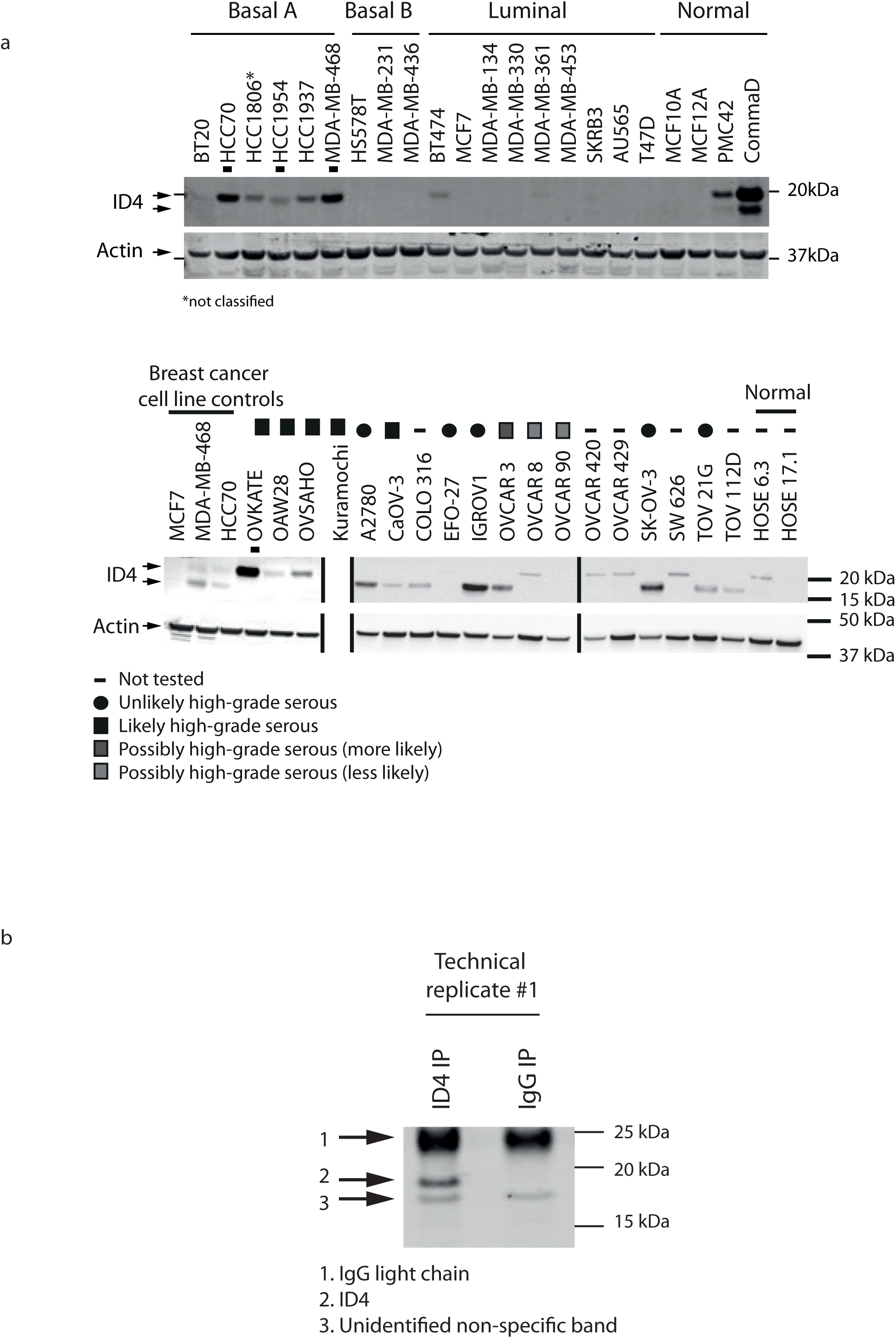
Analysis of ID4 protein expression across a panel of breast and ovarian cell lines. A; Western blotting analysis of ID4 protein expression across a panel of breast (top) and ovarian (bottom) normal and cancer cell lines. A rabbit monoclonal antibody specific to ID4 (Biocheck, BCH-9/82-12) was used for detection (Junankar et al., 2015). Two isoforms of ID4 are detectable across the panel. [β-Actin is shown as a loading control. Modifications to images indicated with vertical black line. Cell lines selected for further analysis are indicated with a black square. Breast cancer subtypes and ovarian cancer histotypes indicated (Neve et al., 2006, Anglesio et al., 2013, Domcke et al., 2013). B; Western blot validation of RIME protocol. MDA-MB-468 cells were processed using the RIME protocol in two technical replicates. Antibodies targeting ID4 (pooled polyclonal antibodies) and IgG (rabbit species matched control) control were used for immunoprecipitation. Western blot analysis for ID4 protein expression, using an independent ID4 monoclonal antibody, following immunoprecipitation. ID4, light chain and non-specific bands labelled.

**Supplementary Figure 2:**
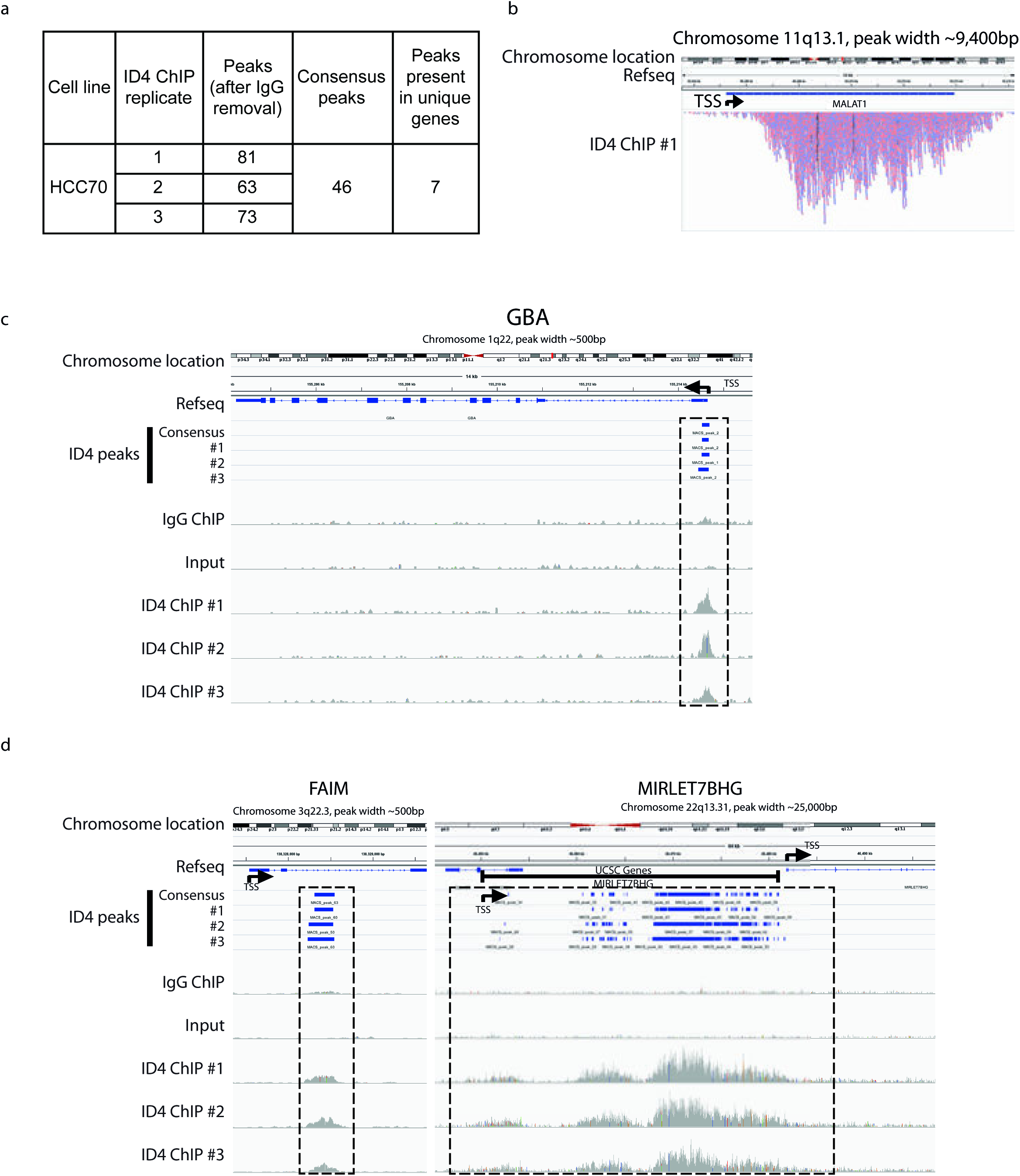
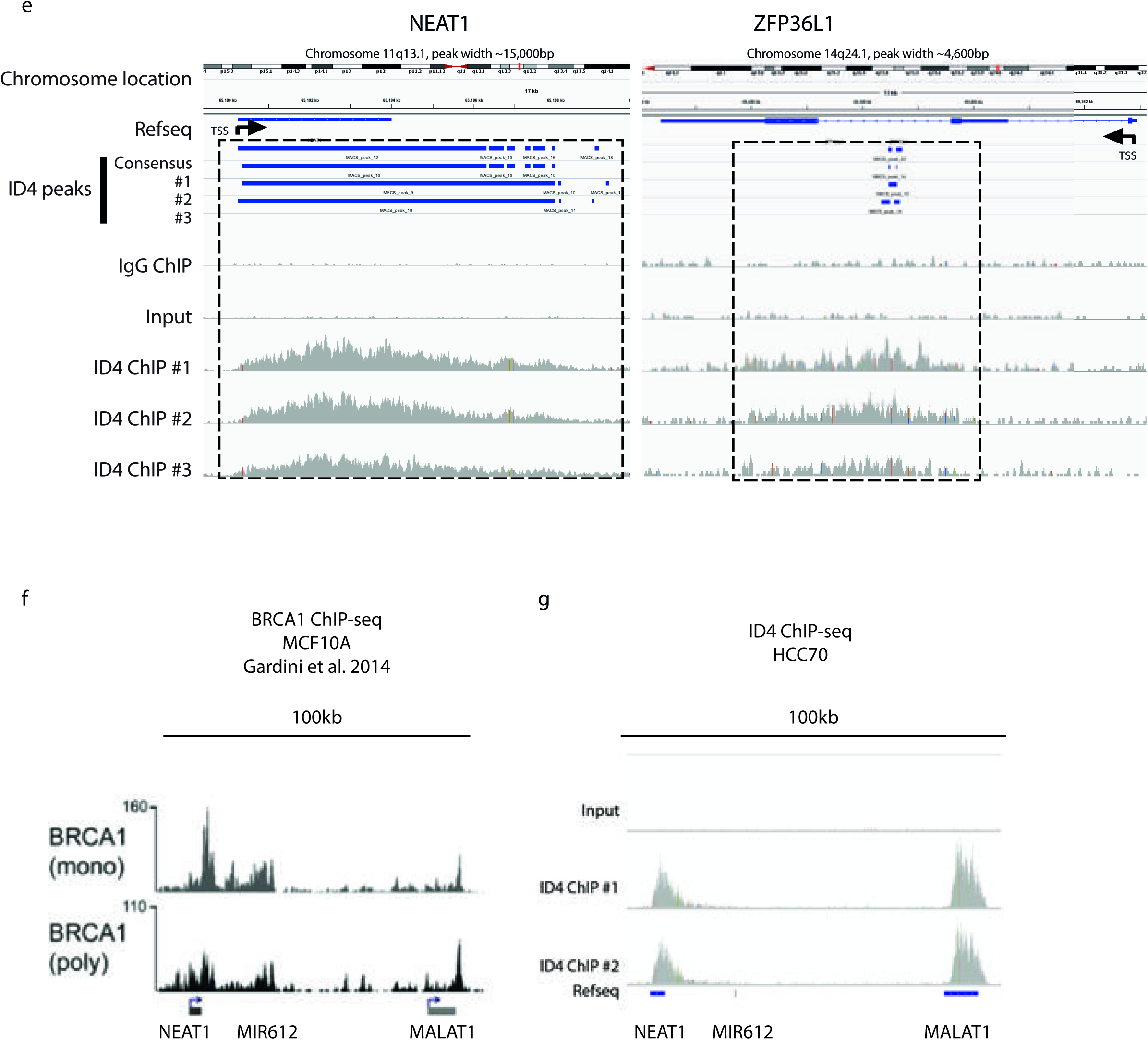
ID4 ChIP-sequencing analysis of HCC70 cell line identifies reproducible ID4-chromatin binding sites. A; Three biological replicates of ID4 ChIP-seq analysis in HCC70 cell line. IgG ChIP-seq and Input controls are shown for comparison. B; 10,000bp resolution image of ID4 ChIP-seq technical replicate #1 binding to MALAT1 gene. Red sequencing reads are aligned to the positive strand (5′ - 3′), and blue to the negative strand of DNA (3′ - 5′). ID4 binding to C; GBA, D; right: FAIM and left: MIRLET7BHG, E; left: NEATland right: ZFP36L1. The chromosomal location, size of the gene and Refseq, human reference genome, are displayed at the top of the image. Reads have been aligned to the human reference genome Hgl9 and peaks called using MACs peak calling algorithm (v2.0.9) (Serandour et al., 2013). Images contain ChIP-seq coverage data and the peaks called for each ID4 technical replicate and the consensus peaks called for all three ID4 ChIP-seq biological replicate for selected gene regions. ID4 binding is shown in comparison to IgG and Input data for the same region. Data visualised using IGV (Thorvaldsdóttir et al., 2013, Robinson et al., 2011). Transcription Start Site (TSS) indicated with black arrow. F; BRCA1 ChIP-seq analysis of NEAT1 and MALAT1 in MCF10A cell line. Displays tracks from two independent BRCA1 antibodies. Modified from (Gardini et al., 2014). G; Input and ID4 ChIP-seq analysis of NEAT1 and MALAT1 genomic loci in HCC70 cell line. Two technical replicates.

**Supplementary Figure 3:**
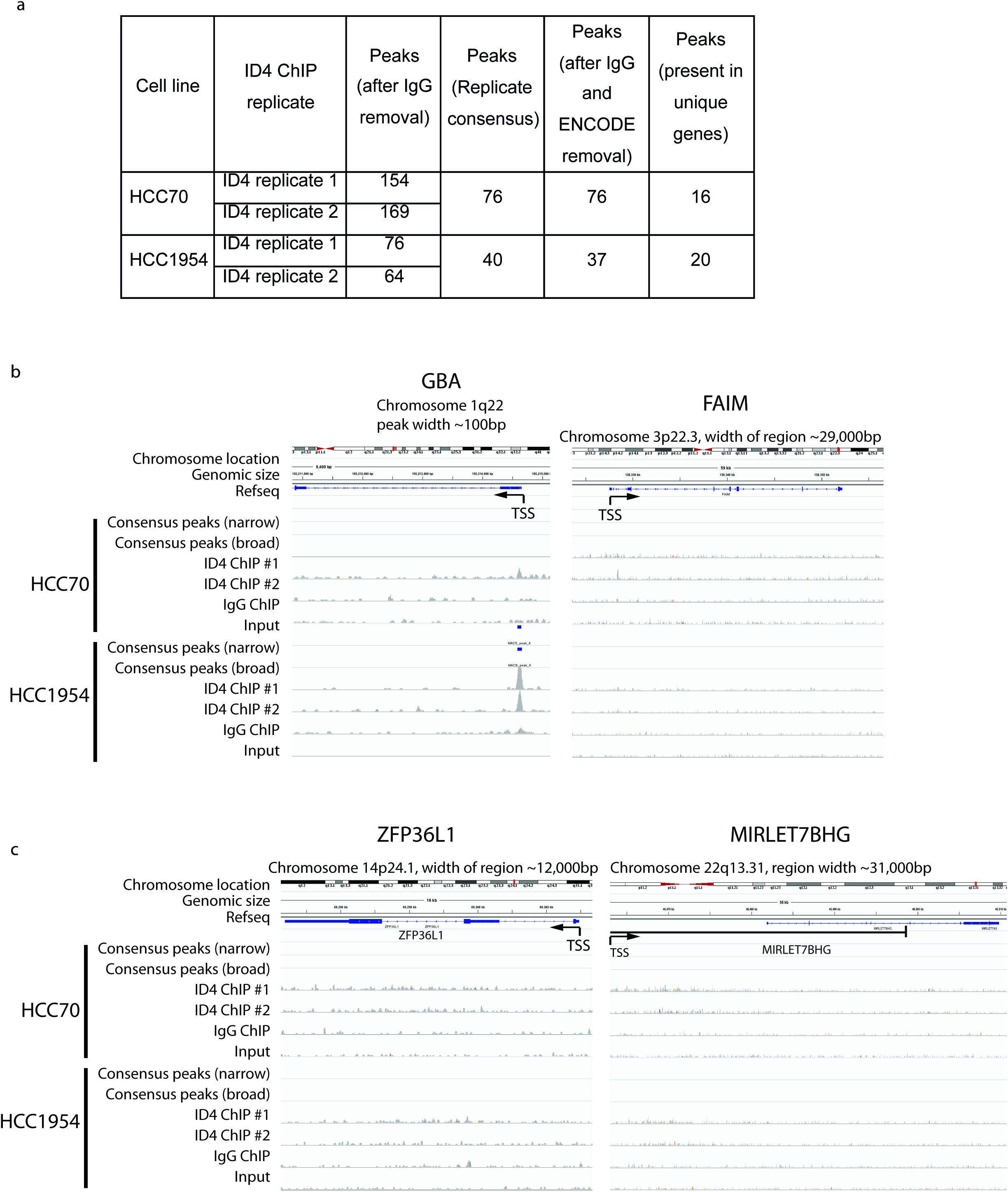
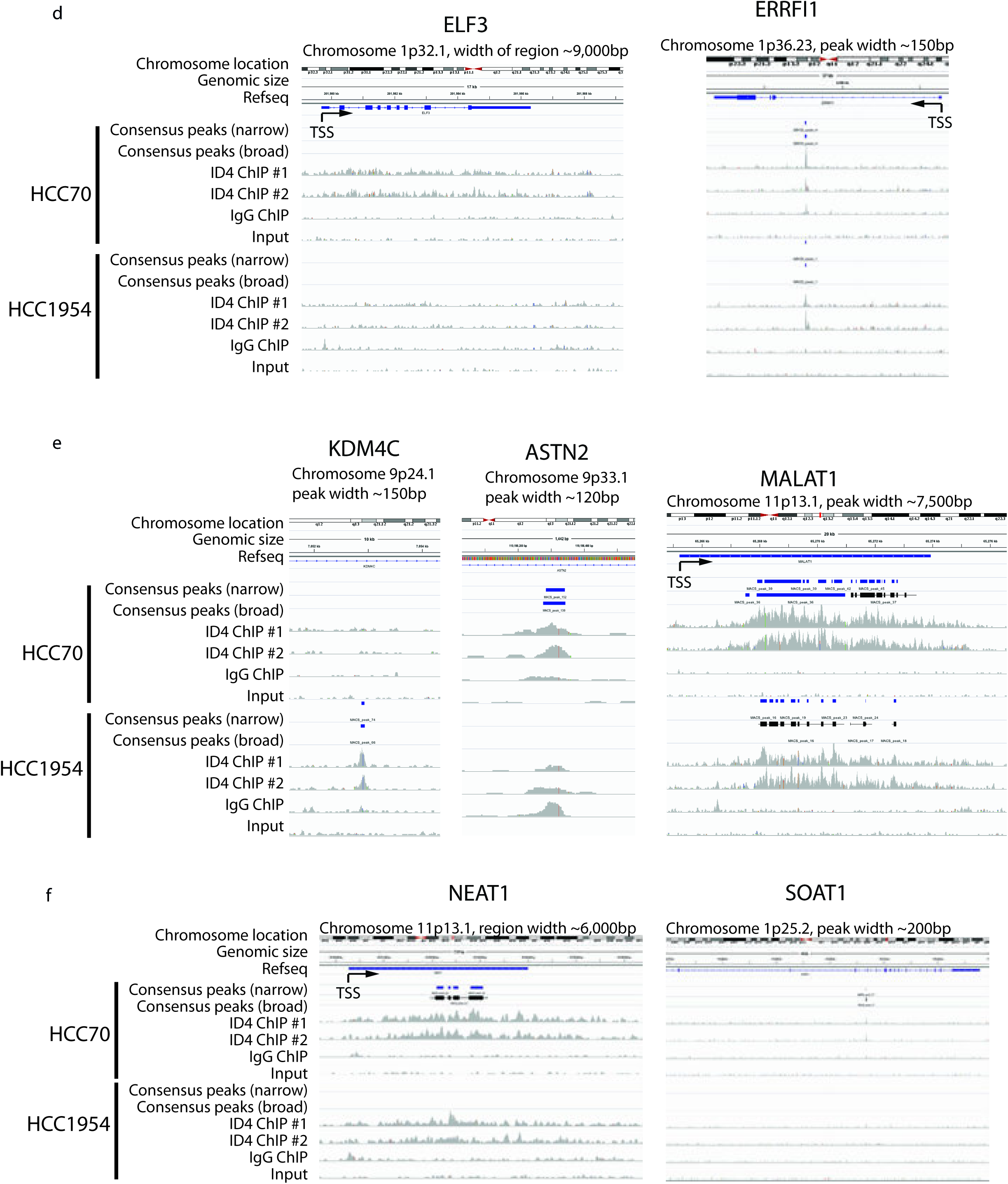
ID4 ChIP-exonuclease sequencing analysis of HCC70 and HCC1954 cell lines reproduces ChIP-seq analysis. A; Table summarising ChIP-exonuclease sequencing analysis of ID4 and IgG binding events in HCC70 and HCC1954 breast cancer cell lines. ID4 binding normalised as for ChIP-seq analysis. B; ChIP-exo analysis of the HCC70 and HCC1954 breast cancer cell lines showing ID4 binding to left: GBA, right: FAIM, C; left: ZFP36L1, right: MIRLET7BHG, D; left: ELF3, right: ERRFI1, E; left: KDMC4, middle: ASTN2, right: SOAT1, F; left: NEAT1, right: MALAT1. Reads have been aligned to the human reference genome Hgl9 and peaks called using MACs peak calling algorithm (v2.0.9) (Serandour et al., 2013). Images contain ChIP-exo coverage data and the peaks called for each ID4 technical replicate and the consensus peaks called for both ID4 ChIP-exo technical replicate for selected gene regions. ID4 binding is shown in comparison to IgG and input data for the same region. Data visualised using IGV (Thorvaldsdottir et al., 2013, Robinson et al., 2011). The chromosomal location and size of the gene are displayed at the top of the image. Below this, Refseq, human reference genome, displays the gene corresponding to particular genomic loci. Transcription Start Site (TSS) indicated with black arrow.

**Supplementary Figure 4:**
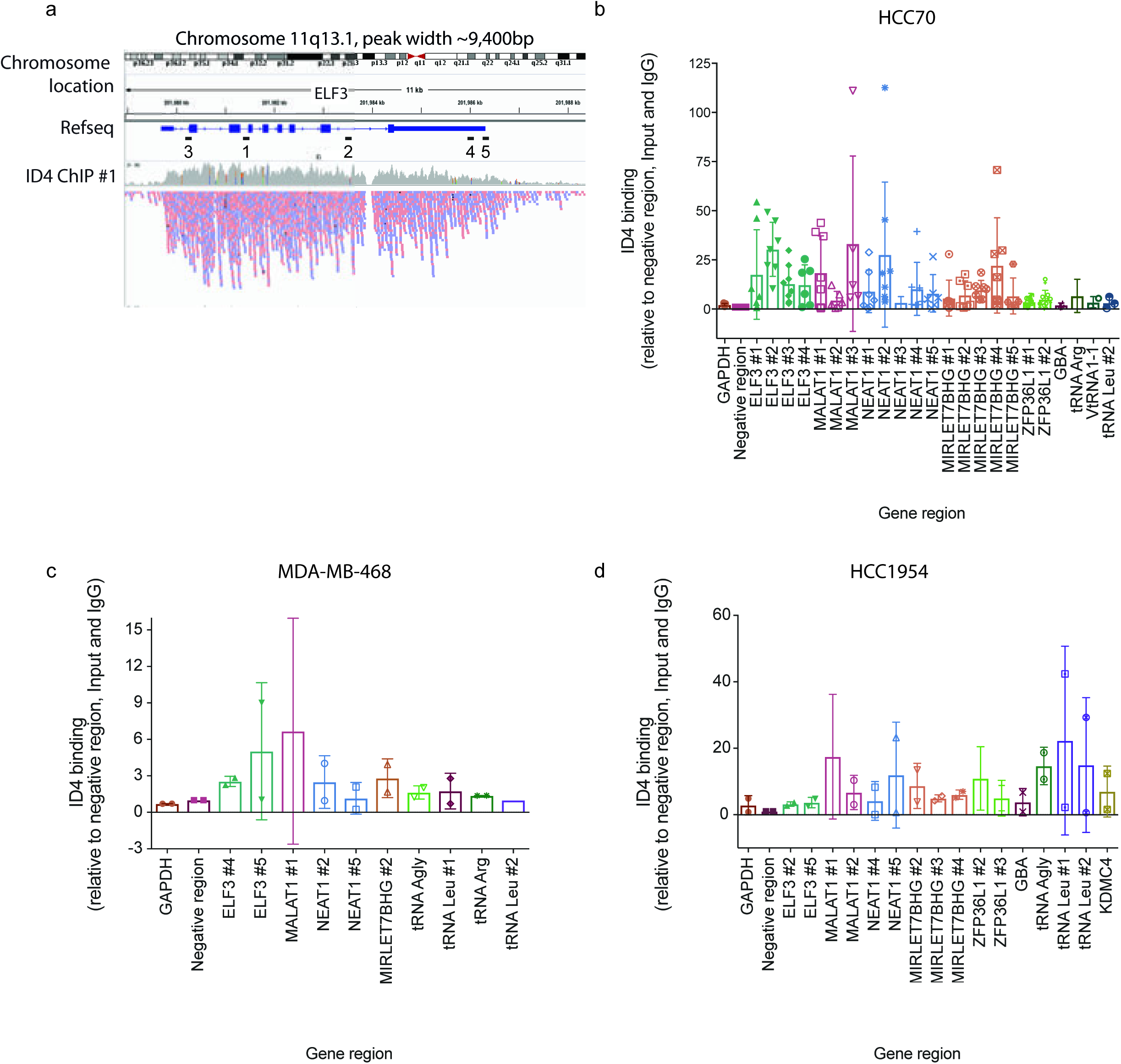
Validation of ID4 binding to specific loci in HCC70, MDA-MB-468 and HCC1954 cell lines. A; Schematic of primer binding across ELF3 gene region. Primers 1-5 are scattered along the length of the ELF3 gene ID4. ChIP-qPCR analysis in B; HCC70 (2-8 biological replicates), C; MDA-MB-468 (2 biological replicates) and D; HCC1954 (2 biological replicates) cells. Multiple primers were designed to tile across the large ID4 binding sites. ID4 binding normalised to input DNA and to a region not bound by ID4 (negative region) and represented as fold-change over IgG control. 1-Way-ANOVA of ID4 binding across all primers compared to negative region: HCC70 p=0.0001, MDA-MB-468 p=0.549, HCC1954 p=0.519.

**Supplementary Figure 5:**
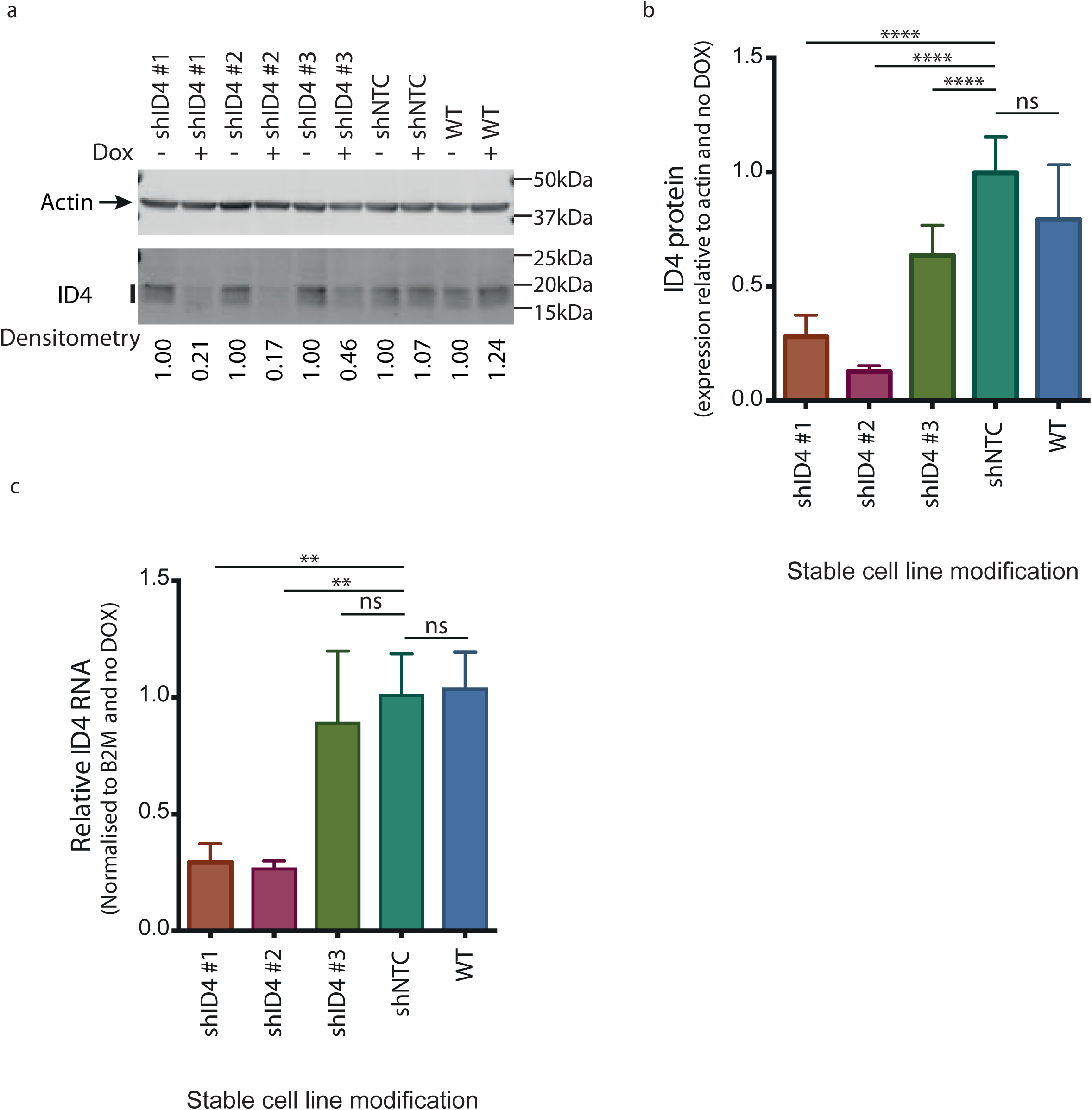
ID4 knockdown using SMARTChoice inducible lentiviral knockdown system. Representative western blot showing ID4 and β-Actin protein expression in wild-type HCC70 cell line and HCC70 cells stably transfected with SMARTChoice vectors containing doxycycline-inducible shRNAs targeting ID4 (shlD4 #1, #2 and #3) or nontargeting control region (shNTC). Cells treated with and without DOX for 72 h. B; Western blotting densitometry quantification of ID4 protein expression across six biological replicates of ID4 knockdown following treatment with and without DOX for 72 h. ID4 expression normalised to β-ACTINand no DOX control. Error bars measure standard error of 4-6 independent replicates. C; qRT-PCR analysis of ID4 RNA expression matched to cells in (B). Data normalised to B2M housekeeping gene and no DOX control. Error bars measure standard error of 4-6 independent replicates. Student′s t-test compares shNTC with the other modified cell lines. ** p<0.01, **** p<0.001.

**Supplementary Figure 6:**
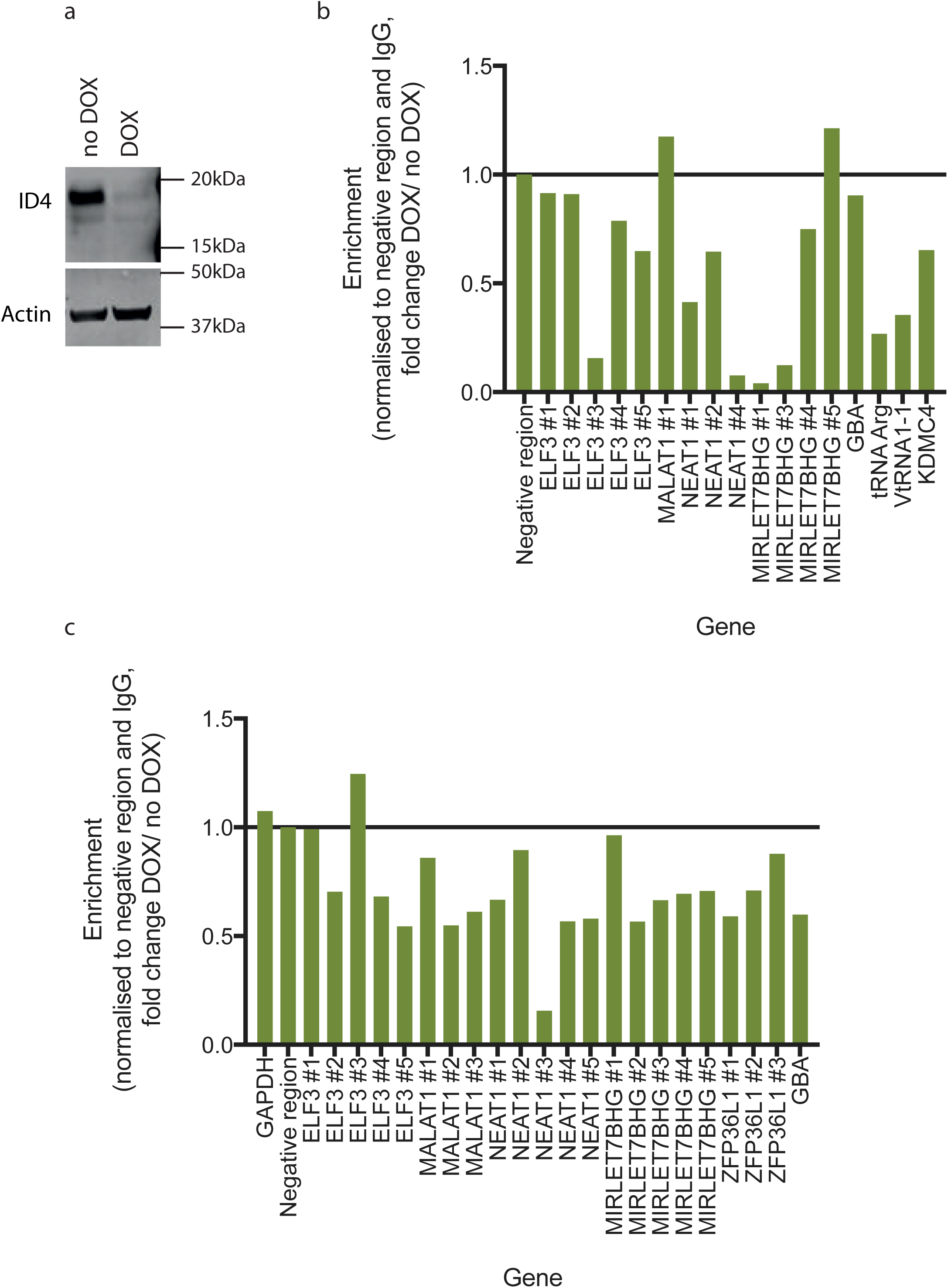
Validation of specificity of ID4-chromatin binding using the SMARTChoice ID4 knockdown model. A; Western blot showing ID4 and β-Actin protein expression in HCC70 cells stably transfected with shlD4 #2 SMARTChoice vector treated with and without doxycycline for 72 h. B; ID4 ChIP-qPCR analysis of cells from (A). ID4 binding normalised to input DNA, to a region not bound by ID4 (negative region) and to the IgG control. Data represented as fold-change over untreated cells. C; replicate of ID4 ChIP-qPCR analysis of HCC70 cells stably transfected with shlD4 #2 SMARTChoice vector treated with and without doxycycline for 72 h.

**Supplementary Figure 7:**
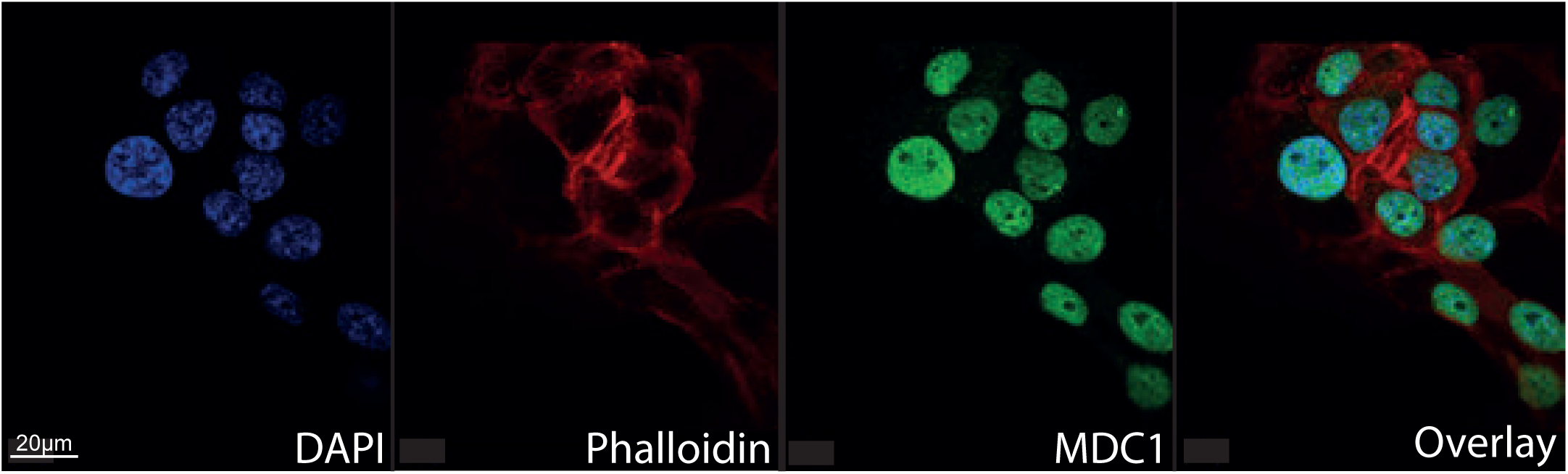
MDC1 is primarily located within the nucleus. Immunofluorescence analysis of MDC1 protein expression in HCC70 cell line. DAPI (nuclear marker); blue, phalloidin (cytoskeletal marker); red and MDC1; green. Imaged using Leica DM 5500 microscope.

**Supplementary Figure 8:**
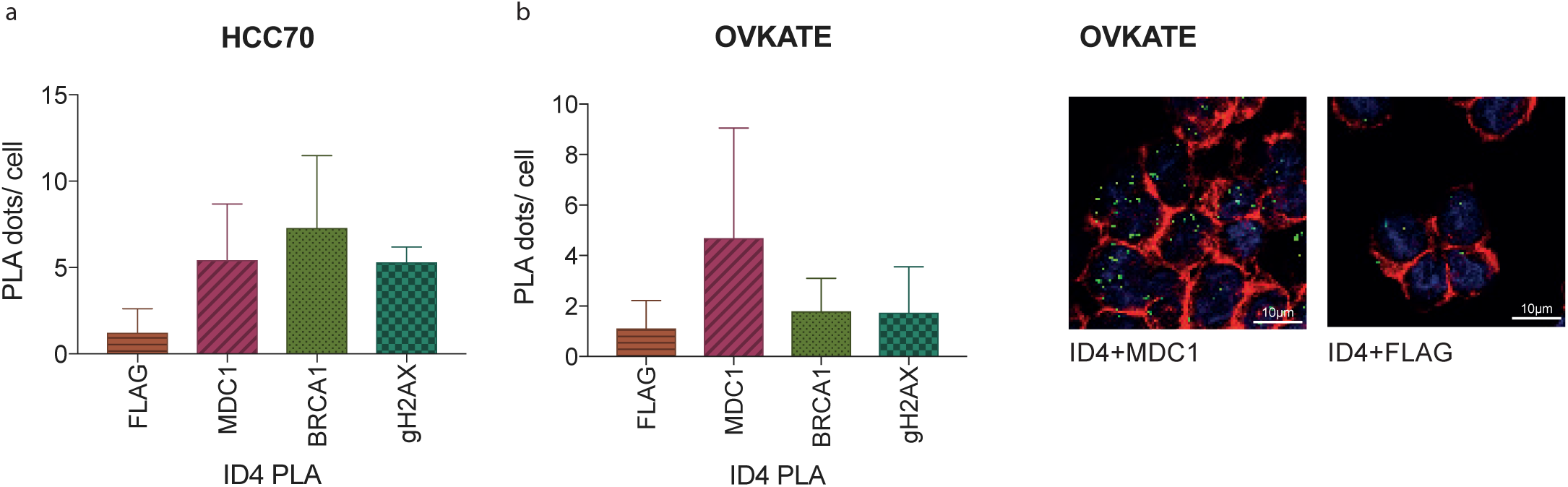
Proximity ligation assay analysis identifies constitutive and DNA damage-inducible binding of ID4 to DNA damage proteins. PLA analysis of ID4 association with FLAG, MDC1, BRCA1 and γH2AX in A; HCC70 and B; OVKATE cell line. Analysis is representative of approximately 150-200 cells of pooled from two independent experiments. 1-way Anova p=0.098. B, right; example image of ID4 and MDC1 PLA. Quantification of interactions presented on right (number of dots/ cell).

**Supplementary Figure 9:**
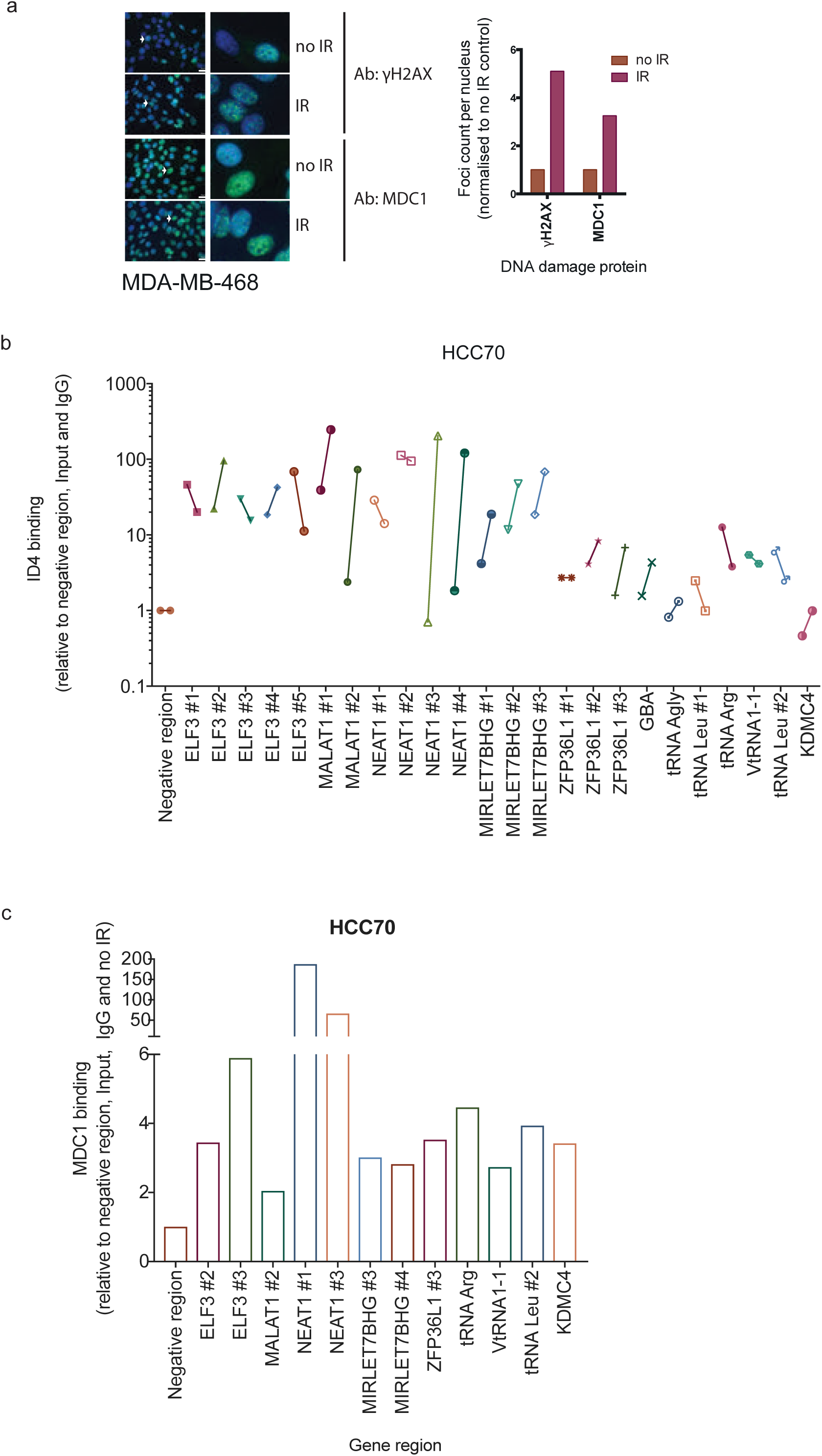
Ionising radiation increases DNA damage foci formation and localisation of MDC1 to chromatin. Induction of DNA damage using ionising radiation measured using immunofluorescence analysis in MDA-MB-468 cells for γH2AX (top) and MDC1 (bottom) foci formation (5Gy with 5 h recovery time prior to fixation). Example images on left, quantification of foci on right. Four to five images were taken for each condition; 50-100 cells. B; Second replicate of HCC70 ID4 ChIP-qPCR. C; ChIP-qPCR analysis of MDC1 positive control binding following ionising radiation DNA damage induction. Data normalised as for Figure 3C, with data shown as fold-change compared to no-IR, n=l.

**Supplementary Figure 10:**
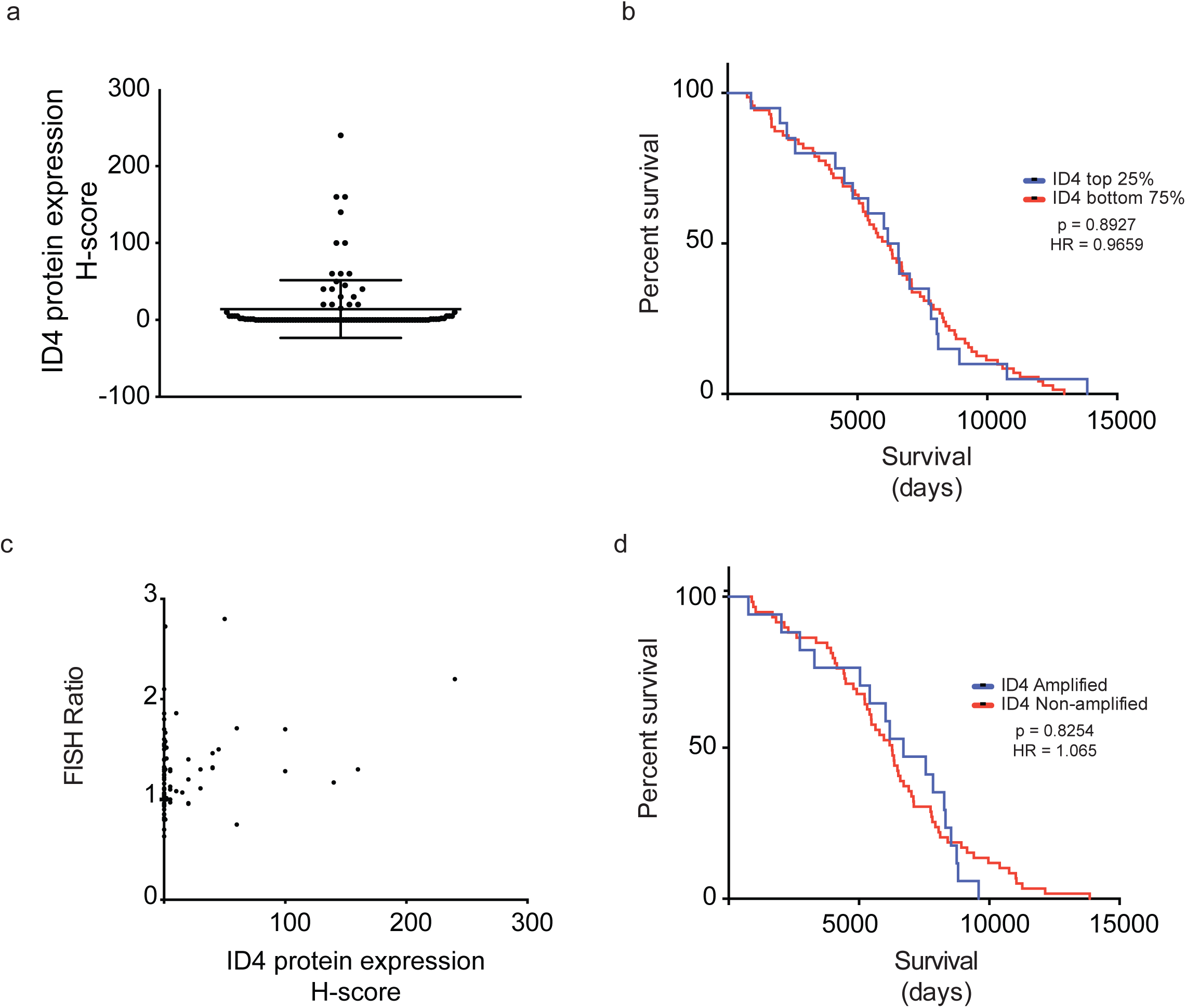
ID4 is overexpressed and amplified at a higher frequency in *BRCA1*-mutant BLBC. A; Graph of ID4 protein expression measured by H-score. Contains 97 *BRCA1*-mutant BLBC patients. B; Survival data for highest 25% of ID4 protein expression (H-score) compared with bottom 75% of ID4 expression (p = 0.8927, HR = 0.9659, 95% Cl = 0.5834 to 1.599). C; ID4 protein expression measured by immunohistochemistry H-score compared with corresponding *ID4:CEP6* FISH ratio. Assuming non-Gaussian distribution, H-score and FISH correlated with a value of r = 0.265 and Spearman correlation p value of <0.00881. D; Survival data for ID4 amplified vs non-amplified cases, (p = 0.825, HR = 1.069, 95% Cl = 0.623 to 1.833).

**Supplementary Figure 11:**
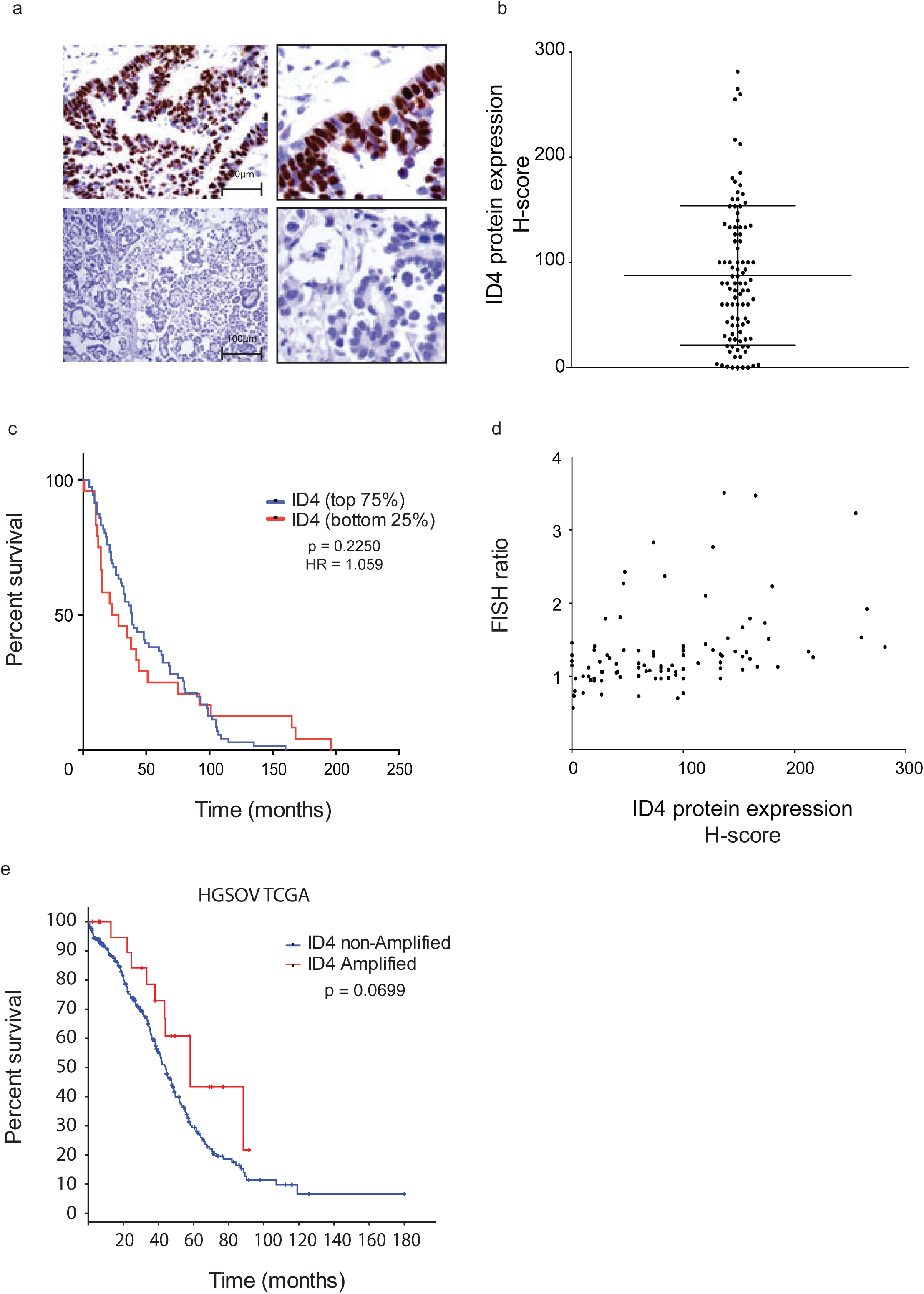
ID4 is overexpressed and amplified in a subset of highgrade serous ovarian cancers. A; Immunohistochemistry staining for ID4 in HGSOV. Example images of ID4 high expressing (top) and ID4 low expressing (bottom) HGSOV. Left, low magnification images, right high magnification. B; Graph of ID4 protein expression measured by H-score. Contains 97 patients. C; Survival data for highest 75% of ID4 protein expression (H-score) compared with bottom 25% of ID4 expression (p = 0.225, HR = 1.059, 95% Cl = 0.646 to 1.735). D; ID4 H-score compared with corresponding *ID4:CEP6* FISH ratio. Assuming non-Gaussian distribution, H-score and FISH correlated with a value of r = 0.457 and Spearman correlation p = <0.0001. E; Analysis of TCGA HGSOV data. Patients divided into *ID4* amplified and *ID4* non-amplified cases. Median survival is 58.1 and 44.1 months, respectively, p = 0.067).

## References

Ando, K., Ozaki, T., Hirota, T. & Nakagawara, A. 2013. NFBD1/MDC1 is phosphorylated by PLK1 and controls G2/M transition through the regulation of a TOPOIIα-mediated decatenation checkpoint. PloS one, 8, e82744.

Andrews, S. 2010. FastQC: A quality control tool for high throughput sequence data. Reference Source.

Anglesio, M. S., Wiegand, K. C., Melnyk, N., Chow, C., Salamanca, C., Prentice, L. M., Senz, J., Yang, W., Spillman, M. A. & Cochrane, D. R. 2013. Type-specific cell line models for type-specific ovarian cancer research. PloS one, 8, e72162.

Asirvatham, A. J., Schmidt, M. A. & Chaudhary, J. 2006. Non-redundant inhibitor of differentiation (Id) gene expression and function in human prostate epithelial cells. The Prostate, 66, 921–935.

Audeh, M. W., Carmichael, J., Penson, R. T., Friedlander, M., Powell, B.,Bell-Mcguinn, K. M., Scott, C., Weitzel, J. N., Oaknin, A. & Loman, N. 2010. Oral poly (ADP-ribose) polymerase inhibitor olaparib in patients with BRCA1 or BRCA2 mutations and recurrent ovarian cancer: a proof-of-concept trial. The Lancet, 376, 245–251.

Beger, C., Pierce, L. N., KrÜGer, M., Marcusson, E. G., Robbins, J. M., Welcsh, P., Welch, P. J., Welte, K., King, M.-C. & Barber, J. R. 2001. Identification of Id4 as a regulator of BRCA1 expression by using a ribozyme-library-based inverse genomics approach. Proceedings of the National Academy of Sciences, 98, 130–135.

Bekker-Jensen, S., Lukas, C., Melander, F., Bartek, J. & Lukas, J. 2005. Dynamic assembly and sustained retention of 53BP1 at the sites of DNA damage are controlled by Mdc1/NFBD1. The Journal of cell biology, 170, 201–211.

Benezra, R., Davis, R. L., Lockshon, D., Turner, D. L. & Weintraub, H. 1990. The protein Id: a negative regulator of helix-loop-helix DNA binding proteins. Cell, 61, 49–59.

Best, S. A., Hutt, K. J., Fu, N. Y., Vaillant, F., Liew, S. H., Hartley, L., Scott, C. L., Lindeman, G. J. & Visvader, J. E. 2014. Dual roles for Id4 in the regulation of estrogen signaling in the mammary gland and ovary. Development, 141, 3159–3164.

Branham, M., Campoy, E., Laurito, S., Branham, R., Urrutia, G., Orozco, J., Gago, F., Urrutia, R. & RoquÉ, M. 2016. Epigenetic regulation of ID4 in the determination of the BRCAness phenotype in breast cancer. Breast cancer research and treatment, 155, 13–23.

Budwit-Novotny, D. A., Mccarty, K. S., Cox, E. B., Soper, J. T., Mutch, D. G.,Creasman, W. T. & Flowers, J. L. 1986. Immunohistochemical analyses of estrogen receptor in endometrial adenocarcinoma using a monoclonal antibody. Cancer research, 46, 5419–5425.

Chaudhary, J., Johnson, J., Kim, G. & Skinner, M. K. 2001. Hormonal Regulation and Differential Actions of the Helix-Loop-Helix Transcriptional Inhibitors of Differentiation (Id1, Id2, Id3, and Id4) in Sertoli Cells 1. Endocrinology, 142, 1727–1736.

Cortez, D., Wang, Y., Qin, J. & Elledge, S. J. 1999. Requirement of ATM-dependent phosphorylation of brca1 in the DNA damage response to double-strand breaks. Science, 286, 1162–1166.

Crippa, E., Lusa, L., De Cecco, L., Marchesi, E., Calin, G. A., Radice, P., Manoukian, S., Peissel, B., Daidone, M. G. & Gariboldi, M. 2014. miR-342 regulates BRCA1 expression through modulation of ID4 in breast cancer. PloS one, 9, e87039.

De Candia, P., Akram, M., Benezra, R. & Brogi, E. 2006. Id4 messenger RNA and estrogen receptor expression: inverse correlation in human normal breast epithelium and carcinoma. Human pathology, 37, 1032–1041.

Derose, Y. S., Gligorich, K. M., Wang, G., Georgelas, A., Bowman, P., Courdy,S. J., Welm, A. L. & Welm, B. E. 2013. Patient-Derived Models of Human Breast Cancer: Protocols for In Vitro and In Vivo Applications in Tumor Biology and Translational Medicine. Current protocols in pharmacology, 14.23. 1–14.23. 43.

Domagala, P., Huzarski, T., Lubinski, J., Gugala, K. & Domagala, W. 2011. PARP-1 expression in breast cancer including BRCA1-associated, triple negative and basal-like tumors: possible implications for PARP-1 inhibitor therapy. Breast cancer research and treatment, 127, 861–869.

Domcke, S., Sinha, R., Levine, D. A., Sander, C. & Schultz, N. 2013. Evaluating cell lines as tumour models by comparison of genomic profiles. Nature communications, 4.

Dong, J., Huang, S., Caikovski, M., Ji, S., Mcgrath, A., Custorio, M. G., Creighton, C. J., Maliakkal, P., Bogoslovskaia, E. & Du, Z. 2011. ID4 regulates mammary gland development by suppressing p38MAPK activity. Development, 138, 5247–5256.

Engel, I. & Murre, C. 2001. The function of E-and Id proteins in lymphocyte development. Nature reviews Immunology, 1, 193.

Ferrer-Vicens, I., Riffo-Campos, Á. L., Zaragozá, R., García, C., López-Rodas, G., Viña, J. R., Torres, L. & García-Trevijano, E. R. 2014. In vivo genome-wide binding of Id2 to E2F4 target genes as part of a reversible program in mice liver. Cellular and Molecular Life Sciences, 71, 3583–3597.

Fontemaggi, G., Dell’Orso, S., Trisciuoglio, D., Shay, T., Melucci, E., Fazi, F., Terrenato, I., Mottolese, M., Muti, P. & Domany, E. 2009. The execution of the transcriptional axis mutant p53, E2F1 and ID4 promotes tumor neo-angiogenesis. Nature structural & molecular biology, 16, 1086–1093.

Furuta, S., Jiang, X., Gu, B., Cheng, E., Chen, P.-L. & Lee, W.-H. 2005. Depletion of BRCA1 impairs differentiation but enhances proliferation of mammary epithelial cells. Proceedings of the National Academy of Sciences of the United States of America, 102, 9176–9181.

Gardini, A., Baillat, D., Cesaroni, M. & Shiekhattar, R. 2014. Genome-wide analysis reveals a role for BRCA1 and PALB2 in transcriptional co-activation. The EMBO journal, e201385567.

Hartman, A. R., Kaldate, R. R., Sailer, L. M., Painter, L., Grier, C. E.,Endsley, R. R., Griffin, M., Hamilton, S. A., Frye, C. A. & Silberman, M. A. 2012. Prevalence of BRCA mutations in an unselected population of triple - negative breast cancer. Cancer, 118, 2787–2795.

Holstege, H., Horlings, H. M., Velds, A., LangerØD, A., BØRresen-Dale, A.-L., Van De Vijver, M. J., Nederlof, P. M. & Jonkers, J. 2010. BRCA1-mutated and basal-like breast cancers have similar aCGH profiles and a high incidence of protein truncating TP53 mutations. BMC cancer, 10, 1.

Huang, Q., Yang, L., Luo, J., Guo, L., Wang, Z., Yang, X., Jin, W., Fang, Y., Ye, J. & Shan, B. 2015. SWATH enables precise label-free quantification on proteome scale. Proteomics, 15, 1215–1223.

Jen, Y., Manova, K. & Benezra, R. 1996. Expression patterns of Id1, Id2, and Id3 are highly related but distinct from that of Id4 during mouse embryogenesis. Developmental Dynamics, 207, 235–252.

Jen, Y., Weintraub, H. & Benezra, R. 1992. Overexpression of Id protein inhibits the muscle differentiation program: in vivo association of Id with E2A proteins. Genes & development, 6, 1466–1479.

Junankar, S., Baker, L. A., Roden, D. L., Nair, R., Elsworth, B., Gallego-Ortega, D., Lacaze, P., Cazet, A., Nikolic, I. & Teo, W. S. 2015. ID4 controls mammary stem cells and marks breast cancers with a stem cell-like phenotype. Nature communications, 6.

Kent, W. J., Sugnet, C. W., Furey, T. S., Roskin, K. M., Pringle, T. H., Zahler, A. M. & Haussler, D. 2002. The human genome browser at UCSC. Genome research, 12, 996–1006.

Konstantinopoulos, P. A., Spentzos, D., Karlan, B. Y., Taniguchi, T.,Fountzilas, E., Francoeur, N., Levine, D. A. & Cannistra, S. A. 2010. Gene expression profile of BRCAness that correlates with responsiveness to chemotherapy and with outcome in patients with epithelial ovarian cancer. Journal of Clinical Oncology, 28, 3555–3561.

Koressaar, T. & Remm, M. 2007. Enhancements and modifications of primer design program Primer3. Bioinformatics, 23, 1289–1291.

Langlands, K., Yin, X., Anand, G. & Prochownik, E. V. 1997. Differential interactions of Id proteins with basic-helix-loop-helix transcription factors. Journal of Biological Chemistry, 272, 19785–19793.

Langmead, B. & Salzberg, S. L. 2012. Fast gapped-read alignment with Bowtie 2. Nature methods, 9, 357.

Law, A. M., Yin, J. X., Castillo, L., Young, A. I., Piggin, C., Rogers, S., Caldon, C. E., Burgess, A., Millar, E. K. & O’Toole, S. A. 2017. Andy’s Algorithms: new automated digital image analysis pipelines for FIJI. Scientific reports, 7, 15717.

Lee, J.-H., Park, S.-J., Hariharasudhan, G., Kim, M.-J., Jung, S. M., Jeong, S.-Y., Chang, I.-Y., Kim, C., Kim, E. & Yu, J. 2017. ID3 regulates the MDC1-mediated DNA damage response in order to maintain genome stability. Nature communications, 8, 903.

Li, H., Handsaker, B., Wysoker, A., Fennell, T., Ruan, J., Homer, N., Marth, G., Abecasis, G. & Durbin, R. 2009. The sequence alignment/map format and SAMtools. Bioinformatics, 25, 2078–2079.

Liebler, D. C. & Zimmerman, L. J. 2013. Targeted quantitation of proteins by mass spectrometry. Biochemistry, 52, 3797–3806.

Lips, E., Mulder, L., Oonk, A., Van Der Kolk, L., Hogervorst, F., Imholz, A., Wesseling, J., Rodenhuis, S. & Nederlof, P. 2013. Triple-negative breast cancer: BRCAness and concordance of clinical features with BRCA1-mutation carriers. British journal of cancer, 108, 2172.

Liu, J., Luo, S., Zhao, H., Liao, J., Li, J., Yang, C., Xu, B., Stern, D. F., Xu, X. & Ye, K. 2012. Structural mechanism of the phosphorylation-dependent dimerization of the MDC1 forkhead-associated domain. Nucleic acids research, 40, 3898–3912.

Liu, L., Zhou, W., Cheng, C.-T., Ren, X., Somlo, G., Fong, M. Y., Chin, A. R., Li, H., Yu, Y. & Xu, Y. 2014. TGFβ induces “BRCAness” and sensitivity to PARP inhibition in breast cancer by regulating DNA-repair genes. Molecular Cancer Research, 12, 1597–1609.

Lou, Z., Minter-Dykhouse, K., Franco, S., Gostissa, M., Rivera, M. A., Celeste, A., Manis, J. P., Van Deursen, J., Nussenzweig, A. & Paull, T. T. 2006. MDC1 maintains genomic stability by participating in the amplification of ATM-dependent DNA damage signals. Molecular cell, 21, 187–200.

Loveys, D. A., Streiff, M. B. & Kato, G. J. 1996. E2A basic-helix-loop-helix transcription factors are negatively regulated by serum growth factors and by the Id3 protein. Nucleic acids research, 24, 2813–2820.

Lu, L. Y., Ou, N. & Lu, Q.-B. 2013. Antioxidant induces DNA damage, cell death and mutagenicity in human lung and skin normal cells. Scientific reports, 3, 3169.

Lukas, C., Melander, F., Stucki, M., Falck, J., Bekker-Jensen, S., Goldberg, M., Lerenthal, Y., Jackson, S. P., Bartek, J. & Lukas, J. 2004. Mdc1 couples DNA double-strand break recognition by Nbs1 with its H2AX-dependent chromatin retention. The EMBO journal, 23, 2674–2683.

Malewicz, M. & Perlmann, T. 2014. Function of transcription factors at DNA lesions in DNA repair. Experimental cell research, 329, 94–100.

Marcotte, R., Sayad, A., Brown, K. R., Sanchez-Garcia, F., Reimand, J.,Haider, M., Virtanen, C., Bradner, J. E., Bader, G. D. & Mills, G. B. 2016. Functional genomic landscape of human breast cancer drivers, vulnerabilities, and resistance. Cell, 164, 293–309.

Martin, M. 2011. Cutadapt removes adapter sequences from high-throughput sequencing reads. EMBnet. journal, 17, pp. 10–12.

Meier, A., Fiegler, H., MuÑOz, P., Ellis, P., Rigler, D., Langford, C., Blasco, M. A., Carter, N. & Jackson, S. P. 2007. Spreading of mammalian DNA-damage response factors studied by ChIP-chip at damaged telomeres. The EMBO journal, 26, 2707–2718.

Melnikova, I. N., Bounpheng, M., Schatteman, G. C., Gilliam, D. & Christy, B. A. 1999. Differential biological activities of mammalian Id proteins in muscle cells. Experimental cell research, 247, 94–104.

Miki, Y., Swensen, J., Shattuck-Eidens, D., Futreal, P. A., Harshman, K.,Tavtigian, S., Liu, Q., Cochran, C., Bennett, L. M. & Ding, W. 1994. A strong candidate for the breast and ovarian cancer susceptibility gene BRCA1. Science, 266, 66–71.

Mohammed, H., D’Santos, C., Serandour, A. A., Ali, H. R., Brown, G. D.,Atkins, A., Rueda, O. M., Holmes, K. A., Theodorou, V. & Robinson, J. L. 2013. Endogenous purification reveals GREB1 as a key estrogen receptor regulatory factor. Cell reports, 3, 342–349.

Montavon, C., Gloss, B. S., Warton, K., Barton, C. A., Statham, A. L., Scurry, J. P., Tabor, B., Nguyen, T. V., Qu, W. & Samimi, G. 2012. Prognostic and diagnostic significance of DNA methylation patterns in high grade serous ovarian cancer. Gynecologic oncology, 124, 582–588.

Mullan, P., Quinn, J. & Harkin, D. 2006. The role of BRCA1 in transcriptional regulation and cell cycle control. Oncogene, 25, 5854.

Natale, F., Rapp, A., Yu, W., Durante, M., Taucher-Scholz, G. & Cardoso, M.C. 2013. Genome-wide multi-parametric analysis of H2AX or γH2AX distributions during ionizing radiation-induced DNA damage response. Epigenetics & Chromatin, 6, 1.

Network, C. G. A. 2012. Comprehensive molecular portraits of human breast tumours. Nature, 490, 61–70.

Network, C. G. A. R. 2011. Integrated genomic analyses of ovarian carcinoma. Nature, 474, 609–615.

Neve, R. M., Chin, K., Fridlyand, J., Yeh, J., Baehner, F. L., Fevr, T., Clark, L., Bayani, N., Coppe, J.-P. & Tong, F. 2006. A collection of breast cancer cell lines for the study of functionally distinct cancer subtypes. Cancer cell, 10, 515–527.

Nguyen, C. T., Gonzales, F. A. & Jones, P. A. 2001. Altered chromatin structure associated with methylation-induced gene silencing in cancer cells: correlation of accessibility, methylation, MeCP2 binding and acetylation. Nucleic acids research, 29, 4598–4606.

Nik-Zainal, S., Davies, H., Staaf, J., Ramakrishna, M., Glodzik, D., Zou, X.,Martincorena, I., Alexandrov, L. B., Martin, S. & Wedge, D. C. 2016. Landscape of somatic mutations in 560 breast cancer whole-genome sequences. Nature, 534, 47–54.

Norton, J. D. 2000. ID helix-loop-helix proteins in cell growth, differentiation and tumorigenesis. Journal of cell science, 113, 3897–3905.

Oliveros, J. C. 2007. VENNY. An interactive tool for comparing lists with Venn Diagrams.

Perk, J., Iavarone, A. & Benezra, R. 2005. Id family of helix-loop-helix proteins in cancer. Nature Reviews Cancer, 5, 603–614.

Perou, C. M., SØRlie, T., Eisen, M. B., Van De Rijn, M., Jeffrey, S. S., Rees, C. A., Pollack, J. R., Ross, D. T., Johnsen, H. & Akslen, L. A. 2000. Molecular portraits of human breast tumours. Nature, 406, 747–752.

Prat, A. & Perou, C. M. 2011. Deconstructing the molecular portraits of breastcancer. Molecular oncology, 5, 5–23.

Pruszko, M., Milano, E., Forcato, M., Donzelli, S., Ganci, F., Di Agostino, S., De Panfilis, S., Fazi, F., Bates, D. O. & Bicciato, S. 2017. The mutant p53 - ID4 complex controls VEGFA isoforms by recruiting lncRNA MALAT1. EMBO reports, 18, 1331-1351.

Pujana, M. A., Han, J.-D. J., Starita, L. M., Stevens, K. N., Tewari, M., Ahn, J. S.,Rennert, G., Moreno, V., Kirchhoff, T. & Gold, B. 2007. Network modeling links breast cancer susceptibility and centrosome dysfunction. Nature genetics, 39, 1338–1349.

Ren, Y., Cheung, H. W., Von Maltzhan, G., Agrawal, A., Cowley, G. S., Weir, B. A., Boehm, J. S., Tamayo, P., Karst, A. M. & Liu, J. F. 2012. Targeted tumor-penetrating siRNA nanocomplexes for credentialing the ovarian cancer oncogene ID4. Science translational medicine, 4, 147ra112- 147ra112.

Roberts, E. C., Deed, R. W., Inoue, T., Norton, J. D. & Sharrocks, A. D. 2001. Id helix-loop-helix proteins antagonize pax transcription factor activity by inhibiting DNA binding. Molecular and cellular biology, 21, 524–533.

Robinson, J. T., ThorvaldsdÓTtir, H., Winckler, W., Guttman, M., Lander, E. S., Getz, G. & Mesirov, J. P. 2011. Integrative genomics viewer. Nature biotechnology, 29, 24–26.

RodrÍGuez, J. L., Sandoval, J., Serviddio, G., Sastre, J., Morante, M., Perrelli, M.-G., MartÍNez-Chantar, M. L., ViÑA, J., ViÑA, J. R. & Mato, J. M. 2006. Id2 leaves the chromatin of the E2F4–p130-controlled c-myc promoter during hepatocyte priming for liver regeneration. Biochemical Journal, 398, 431–437.

RoldÁN, G., Delgado, L. & MusÉ, I. M. 2006. Tumoral expression of BRCA1, estrogen receptor alpha and ID4 protein in patients with sporadic breast cancer. Cancer biology & therapy, 5, 505–510.

Rouzier, R., Perou, C. M., Symmans, W. F., Ibrahim, N., Cristofanilli, M., Anderson, K., Hess, K. R., Stec, J., Ayers, M. & Wagner, P. 2005. Breast cancer molecular subtypes respond differently to preoperative chemotherapy. Clinical Cancer Research, 11, 5678–5685.

Ruzinova, M. B. & Benezra, R. 2003. Id proteins in development, cell cycle and cancer. Trends in cell biology, 13, 410–418.

Schefe, J. H., Lehmann, K. E., Buschmann, I. R., Unger, T. & Funke-Kaiser, H. 2006. Quantitative real-time RT-PCR data analysis: current concepts and the novel “gene expression’s C T difference” formula. Journal of molecular medicine, 84, 901–910.

Schindelin, J., Arganda-Carreras, I., Frise, E., Kaynig, V., Longair, M., Pietzsch, T., Preibisch, S., Rueden, C., Saalfeld, S. & Schmid, B. 2012. Fiji: an open-source platform for biological-image analysis. Nature methods, 9, 676–682.

Schmidt, D., Wilson, M. D., Spyrou, C., Brown, G. D., Hadfield, J. & Odom, D.T. 2009. ChIP-seq: using high-throughput sequencing to discover protein–DNA interactions. Methods, 48, 240–248.

Scully, R., Chen, J., Ochs, R. L., Keegan, K., Hoekstra, M., Feunteun, J. & Livingston, D. M. 1997. Dynamic changes of BRCA1 subnuclear location and phosphorylation state are initiated by DNA damage. Cell, 90, 425–435.

Scully, R., Xie, A. & Nagaraju, G. 2004. Molecular functions of BRCA1 in the DNA damage response. Cancer biology & therapy, 3, 521–527.

Seo, J., Kim, S. C., Lee, H.-S., Kim, J. K., Shon, H. J., Salleh, N. L. M., Desai, K. V., Lee, J. H., Kang, E.-S. & Kim, J. S. 2012. Genome-wide profiles of H2AX and γ-H2AX differentiate endogenous and exogenous DNA damage hotspots in human cells. Nucleic acids research, gks287.

Serandour, A. A., Brown, G. D., Cohen, J. D. & Carroll, J. S. 2013. Development of an Illumina-based ChIP-exonuclease method provides insight into FoxA1-DNA binding properties. Genome biology, 14, 1.

Shan, L., Yu, M., Qiu, C. & Snyderwine, E. G. 2003. Id4 regulates mammary epithelial cell growth and differentiation and is overexpressed in rat mammary gland carcinomas. The American journal of pathology, 163, 2495–2502.

Shi, W., Ma, Z., Willers, H., Akhtar, K., Scott, S. P., Zhang, J., Powell, S. & Zhang, J. 2008. Disassembly of MDC1 foci is controlled by ubiquitin-proteasome-dependent degradation. Journal of Biological Chemistry, 283, 31608–31616.

SØRlie, T., Perou, C. M., Tibshirani, R., Aas, T., Geisler, S., Johnsen, H., Hastie, T., Eisen, M. B., Van Derijn, M. & Jeffrey, S. S. 2001. Gene expression patterns of breast carcinomas distinguish tumor subclasses with clinical implications. Proceedings of the National Academy of Sciences, 98, 10869–10874.

Stewart, G. S., Wang, B., Bignell, C. R., Taylor, A. M. R. & Elledge, S. J. 2003. MDC1 is a mediator of the mammalian DNA damage checkpoint. Nature, 421, 961–966.

Stucki, M., Clapperton, J. A., Mohammad, D., Yaffe, M. B., Smerdon, S. J. & Jackson, S. P. 2005. MDC1 directly binds phosphorylated histone H2AX to regulate cellular responses to DNA double-strand breaks. Cell, 123, 1213–1226.

Taberlay, P. C., Kelly, T. K., Liu, C.-C., You, J. S., De Carvalho, D. D., Miranda, T. B., Zhou, X. J., Liang, G. & Jones, P. A. 2011. Polycomb-repressed genes have permissive enhancers that initiate reprogramming. Cell, 147, 1283–1294.

Takenaka, K., Nakagawa, H., Miyamoto, S. & Hiroaki, M. 2004. The pre-mRNA-splicing factor SF3a66 functions as a microtubule-binding and- bundling protein. Biochemical Journal, 382, 223–230.

Tassone, P., Di Martino, M. T., Ventura, M., Pietragalla, A., Cucinotto, I., Calimeri, T., Neri, P., Caraglia, M., Tagliaferri, P. & Bulotta, A. 2009. Loss of BRCA1 function increases the antitumor activity of cisplatin against human breast cancer xenografts in vivo. Cancer biology & therapy, 8, 648–653.

Thike, A. A., Tan, P. H., Ikeda, M. & Iqbal, J. 2015. Increased ID4 expression, accompanied by mutant p53 accumulation and loss of BRCA1/2 proteins in triple-negative breast cancer, adversely affects survival. Histopathology.

ThorvaldsdÓTtir, H., Robinson, J. T. & Mesirov, J. P. 2013. Integrative Genomics Viewer (IGV): high-performance genomics data visualization and exploration. Briefings in bioinformatics, 14, 178–192.

Townsend, K., Mason, H., Blackford, A. N., Miller, E. S., Chapman, J. R., Sedgwick, G. G., Barone, G., Turnell, A. S. & Stewart, G. S. 2009. Mediator of DNA damage checkpoint 1 (MDC1) regulates mitotic progression. Journal of Biological Chemistry, 284, 33939-33948.

Turner, N. & Reis-Filho, J. 2006. Basal-like breast cancer and the BRCA1 phenotype. Oncogene, 25, 5846–5853.

Turner, N., Reis-Filho, J., Russell, A., Springall, R., Ryder, K., Steele, D., Savage, K., Gillett, C., Schmitt, F. & Ashworth, A. 2007. BRCA1 dysfunction in sporadic basal-like breast cancer. Oncogene, 26, 2126–2132.

Turner, N., Tutt, A. & Ashworth, A. 2004. Hallmarks of’BRCAness’ in sporadic cancers. Nature Reviews Cancer, 4, 814–819.

Untergasser, A., Cutcutache, I., Koressaar, T., Ye, J., Faircloth, B. C., Remm, M. & Rozen, S. G. 2012. Primer3—new capabilities and interfaces. Nucleic acids research, 40, e115–e115.

Vafaee, F., Colvin, E. K., Mok, S. C., Howell, V. M. & Samimi, G. 2017. Functional prediction of long non-coding RNAs in ovarian cancer-associated fibroblasts indicate a potential role in metastasis. Scientific reports, 7, 10374.

Vollebergh, M., Lips, E., Nederlof, P., Wessels, L., Schmidt, M., Van Beers, E., Cornelissen, S., Holtkamp, M., Froklage, F. & De Vries, E. 2010. An aCGH classifier derived from BRCA1-mutated breast cancer and benefit of high-dose platinum-based chemotherapy in HER2-negative breast cancer patients. Annals of Oncology, 22, 1561–1570.

Wen, Y. H., Ho, A., Patil, S., Akram, M., Catalano, J., Eaton, A., Norton, L.,Benezra, R. & Brogi, E. 2012. Id4 protein is highly expressed in triple-negative breast carcinomas: possible implications for BRCA1 downregulation. Breast cancer research and treatment, 135, 93–102.

Wilhelmsen, K., Litjens, S. H., Kuikman, I., Tshimbalanga, N., Janssen, H.,Van Den Bout, I., Raymond, K. & Sonnenberg, A. 2005. Nesprin-3, a novel outer nuclear membrane protein, associates with the cytoskeletal linker protein plectin. The Journal of cell biology, 171, 799–810.

Wilson, K. A., Colavito, S. A., Schulz, V., Wakefield, P. H., Sessa, W., Tuck, D. & Stern, D. F. 2011. NFBD1/MDC1 regulates Cav1 and Cav2 independently of DNA damage and p53. Molecular Cancer Research, 9, 766–781.

Wu, J., Lu, L.-Y. & Yu, X. 2010. The role of BRCA1 in DNA damage response. Protein & cell, 1, 117–123.

Zhang, F., Bick, G., Park, J.-Y. & Andreassen, P. R. 2012. MDC1 and RNF8 function in a pathway that directs BRCA1-dependent localization of PALB2 required for homologous recombination. J Cell Sci, 125, 6049–6057.

Zhang, Y., Liu, T., Meyer, C. A., Eeckhoute, J., Johnson, D. S., Bernstein, B. E., Nusbaum, C., Myers, R. M., Brown, M. & Li, W. 2008. Model-based analysis of ChIP-Seq (MACS). Genome biology, 9, R137.

